# A near-complete chromosome from a bacterial pathogen integrated into the genome of its arthropod vector

**DOI:** 10.64898/2026.09.16.751987

**Authors:** Jing Jing Khoo, Alexandra Beliavskaia, Adriana Ludwig, Mark Whitehead, Alaa M. Al-Khafaji, Catherine S. Hartley, Maria Kazimirova, Germanus S. Bah, Andeliza Frederik, Inswasti Cahyani, Matthew Loose, Lesley Bell-Sakyi, Alistair C. Darby, Benjamin L. Makepeace

## Abstract

Lateral gene transfer (LGT) from organellar to eukaryotic genomes is ubiquitous, resulting in the presence of widespread nuclear mitochondrial DNA segments (NUMTs) and/or plastid gene transfers. While LGT from bacterial associates of eukaryotes is generally less common, many arthropod genomes are littered with LGT originating from *Wolbachia* (order Rickettsiales) - a vertically transmitted, obligate intracellular symbiont of invertebrates. This contrasts with *Rickettsia*, a related genus including many important human pathogens, which has not been shown to promulgate LGT within its arthropod vectors. Here, we present the 8.6 Gb genome of a cell line derived from *Amblyomma variegatum* (the tropical bont tick), which is the primary vector of *Rickettsia africae* (agent of African tick-bite fever). In addition to numerous NUMTs and diverse families of transposable elements, an almost-complete chromosome of *R. africae* origin was identified in the cell line genome, localised to the putative sex chromosome. Sequencing of field-collected *A. variegatum* confirmed the presence of this large LGT in wild ticks, although the absence of transferred genes from the *R. africae* plasmid provided a means to differentiate LGT from genuine rickettsial infections. These findings highlight the imperative to consider the possibility of LGT when screening vectors for pathogens by PCR.

## Introduction

Endosymbiotic gene transfer from organellar genomes of prokaryotic origin to eukaryotic host genomes is a key feature of most eukaryotic taxa and has been hypothesised to be driven by energy efficiency^1^. Nuclear mitochondrial DNA segments (NUMTs) and nuclear plastid gene transfers (NUPTs) are frequent in eukaryotic genomes, leading to the accumulation of a much greater number of pseudogenes compared to functional gene transfers^2,3^. Crucially, the latter must acquire a nuclear promoter for expression and a targeting motif in the coding sequences (CDS) for successful export of the encoded protein to the organelle of origin. While NUMTs and NUPTs are highly prevalent in eukaryotes, lateral gene transfer (LGT) from non-organellar prokaryotes into eukaryotic genomes, while more common than previously assumed^4,5^, is by no means ubiquitous – particularly for LGT conferring novel functions. However, in the Arthropoda, LGT has played an essential role in diversification of dietary sources and defence against predators or chemical toxins. Functional LGT of bacterial origin into insect genomes is especially important in phytophagous arthropods, such as certain Hemiptera and Lepidoptera, and in the Acari^6,7^. Genes encoding bacterial enzymes transferred into arthropod genomes have been shown to detoxify pesticides^8^ or plant-derived defensive toxins^7^, or to provision vitamins deficient in sap diets^9^. In rarer cases, LGT can also provide arthropods with novel venom components^10^ or other toxins for predation or defence^11^.

Despite these examples of evolutionary innovations conferred by bacterial LGT in Arthropoda, the vast majority of LGT in arthropod genomes appears to be nonfunctional. The largest single source of bacterial LGT in the Arthropoda is from an obligate intracellular, vertically-transmitted symbiont, *Wolbachia* (order Rickettsiales^12^), which infects >50% of terrestrial arthropods^13^. In one extreme example, multiple copies of the *Wolbachia* symbiont chromosome were found to be integrated into the genome of *Drosophila ananassae*^14^, but these insertions are mostly transcriptionally silent^15^. Across the Arthropoda, *Wolbachia* LGTs are generally pseudogenized, although there is one case where LGT from a *Wolbachia* strain that can feminise male isopods recapitulates the same phenotype in the absence of *Wolbachia* infection^16^, suggesting that one or more symbiont genes are functional in their new genomic context. However, the general pattern of pseudogenisation of *Wolbachia* LGT has provided some confidence that live infections can be discriminated from LGT events^17^, even when using PCR assays rather than whole genome sequencing to detect the symbiont^18^.

While *Wolbachia* cannot infect vertebrates, other Rickettsiales such as *Rickettsia* spp. include several major human pathogens. In common with *Wolbachia*, *Rickettsia* is vertically transmitted in the arthropod host (often a disease vector) via the maternal line to the next generation. Surprisingly, LGT of rickettsial genes into the genomes of arthropod hosts appears to be rare, despite the high prevalence of symbiosis involving *Rickettsia*^13^ or its recently described sister taxon, *Candidatus* Tisiphia^19^.

Here, we demonstrate that the genome of the tropical bont tick (*Amblyomma variegatum*) contains LGT spanning almost the entire chromosome of *Rickettsia africae*, the agent of a zoonotic febrile illness termed African tick bite fever (ATBF)^20^. This LGT, which we show is highly likely to be nonfunctional, was first suspected due to detection of *R. africae* gene sequences in two *A. variegatum* cell lines in which no bacteria were visible^21^. Since *A. variegatum* is the primary vector of *R. africae*^22^, this discovery has important implications for epidemiological analyses of ATBF and has wider ramifications for PCR-based studies of pathogens in arthropod hosts for which high-quality genomes are not yet available.

## Results

### A high-quality assembly reveals the massive genome of an *A. variegatum*-derived cell line

To investigate the preliminary discovery of *R. africae* gene sequences in the cell lines derived from *A. variegatum* cells^21^, we sequenced the genome of one of the two cell lines, AVL/CTVM17, on the PacBio, ONT and Illumina platforms. Due to higher contiguity and BUSCO (Supplementary Table S1), the PacBio assembly was combined with a Hi-C library from the same cell line to generate a scaffold-level assembly comprising 721 contigs, with an N_50_ of 1.02 Gb and a total assembly size of ∼8.61 Gb. The final assembly had a BUSCO score of 93.3% (77.6% single copy genes, 15.7% duplicated genes, 3.4% fragmented genes, 3.3% missing genes), indicating a high level of completeness. The chromosome architecture of the cell line resembles that of the source tick species, as shown by the contact heatmap from the cell line assembly and a Hi-C library generated from *A. variegatum* ticks from a colony maintained at the Slovak Academy of Sciences (Fig. 1A) (Supplementary Table S1). Ticks of the genus *Amblyomma*, including *A. variegatum*, are commonly diploid with 20 autosomes and an XX:X0 sex determination system^23,24^. In contrast, the cell line assembly comprised ten major scaffolds (∼457 Mb to ∼1.3 Gb), together accounting for more than 99% of the assembled genome (Fig. 1B), suggesting either that a small chromosome was not represented in the assembly or two chromosomes are fused. We also sequenced field-collected *A. variegatum* ticks from Cameroon on the Illumina platform (Supplementary Methods, Table S1). Mapping of the Illumina reads generated from female (n = 6) and male (n = 9) ticks to the assembly indicated that the second largest scaffold is the X chromosome, as it had consistently lower coverage in male ticks (Fig. 1C). Karyotyping of AVL/CTVM17 cells revealed a modal chromosome number of 42 (*i.e*., multiples of 2n = 20 + X chromosomes) and a range of 6-77 chromosomes per cell (Supplementary Fig. S1). This is consistent with reports of changes in chromosome numbers in other tick cell lines, which could be explained by chromosomal duplication or fusion^25^.

**Fig. 1.**
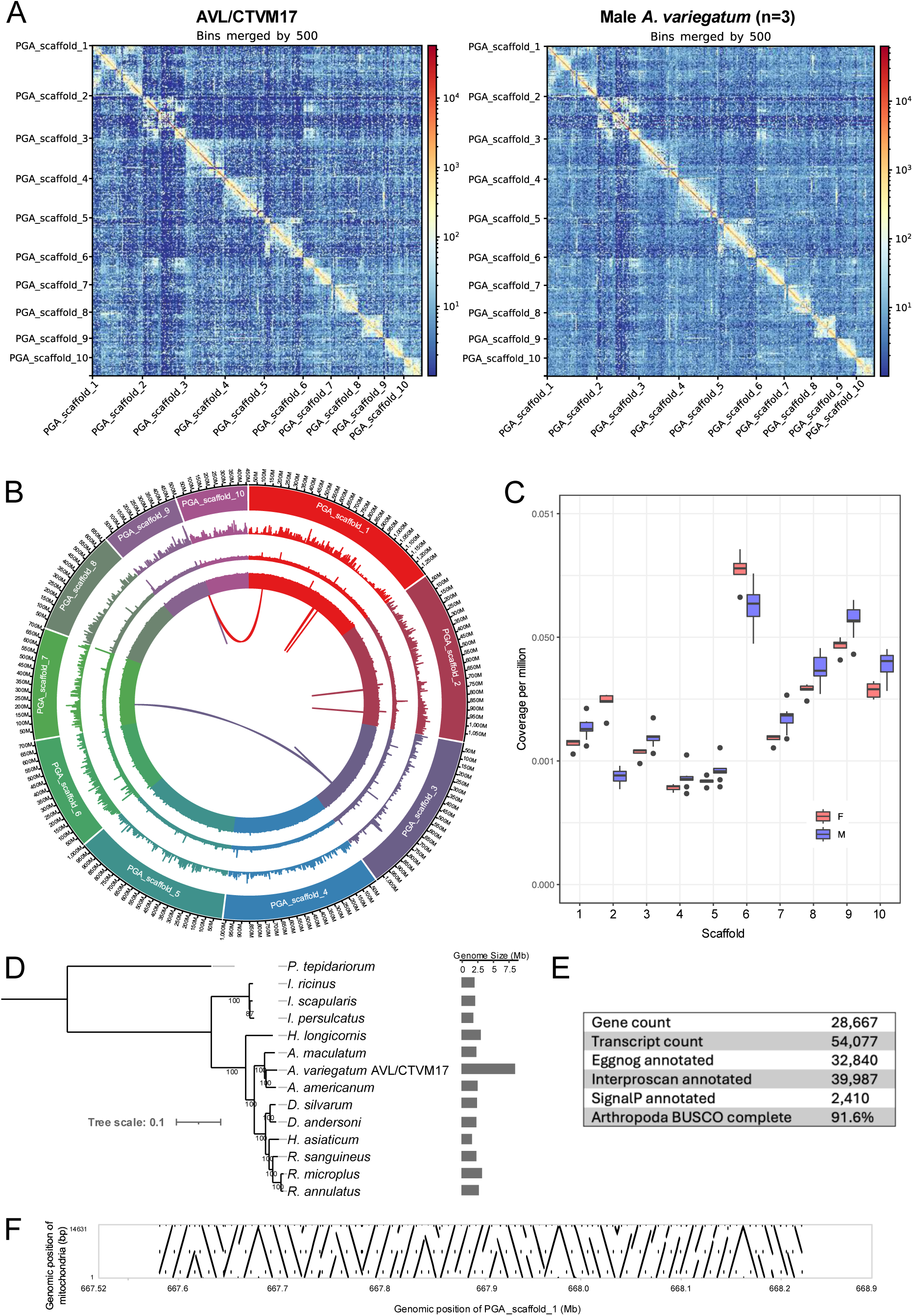
Scaffold level genome assembly from *Amblyomma variegatum* cell line AVL/CTVM17. (A) Contact heatmap using the Hi-C library from the cell line or a pool of three male colony ticks. (B) Circos plot of the ten largest scaffolds. Information is depicted as the following: Outer ring, scaffold lengths in bp (100 Kb sliding window); first inner ring, gene density; second inner ring, repeat density. (C) Illumina sequence coverage across the ten largest scaffolds in female (n = 5) and male (n = 9) ticks. (D) Maximum likelihood tree inferred from 289 shared single-copy orthologues, using the JTT+I+G4 model and 1000 bootstrap replicates. The common house spider, *Parasteatoda tepidariorum*, was used as the outgroup. (E) Genome annotation statistics. (F) Alignment of the *A. variegatum* mitochondrial sequence to PGA_scaffold_1, showing the largest region of NUMTs in the Hi-C assembly.

A phylogenetic tree generated from 289 shared single-copy orthologues across thirteen tick species available at time of study showed that the current assembly is positioned in a clade with two recently sequenced New World *Amblyomma* species, *Amblyomma americanum* (lone star tick) and *Amblyomma maculatum* (Gulf Coast tick) (Fig. 1D). At ∼8.61 Gb, the assembly from AVL/CVTM17 represents the largest tick genome sequenced to date (Fig. 1D). However, whole-genome alignment of the final assembly with the assemblies for *A. americanum* and *A. maculatum* showed little alignment and synteny across the scaffolds (Supplementary Fig. S2).

### Genome annotation and collinearity analysis reveals no evidence of large segmental duplications

In total, 28,667 protein-coding genes and 54,077 potential transcripts were predicted in the current AVL/CTVM17 Hi-C assembly (Fig. 1E). The BUSCO analysis of the protein dataset showed 91.6% completeness (74.3% single copy genes, 17.3% duplicated genes, 1.5% fragmented genes, 6.9% missing genes). EggNOG, InterProScan, and SignalP annotations were also obtained for 32,840, 39,987 and 2,410 transcripts, respectively. Approximately 5.83 Gb, or 68.4% of the assembly, was identified as repetitive DNA (Fig. 1B), potentially containing bona-fide transposable elements (TEs) (see annotation results below) and other repetitive sequences.

Collinearity analysis among the ten largest scaffolds to detect large segmental duplications, identified only 25 colinear gene blocks encompassing 216 genes in total (Supplementary Table S2). The colinear gene blocks mostly occurred within the same scaffold, occasionally in tandem, except for two blocks that were detected in scaffolds 1-9 and scaffolds 3-7 (Fig. 1B). The size of the scaffold regions containing colinear gene blocks ranged from ∼30 Kb to 2.4 Mb (Supplementary Table S2), representing only a tiny fraction of the scaffold lengths (<1%), suggesting that duplication of large genome segments is unlikely to explain the large assembly size (Fig. 1D).

### The AVL/CTVM17 assembly contains a large array of NUMTs

During our earlier attempts to assemble the mitochondrial sequence from the AVL/CTVM17 cell line^26^, multiple mitogenomic variants were obtained from the MitoHifi assembly pipeline (Supplementary Fig. S3), suggesting the presence of NUMTs. We aligned the mitochondrial assembly (14,631 bp) to the Hi-C scaffolds and identified NUMTs in eight of the ten largest scaffolds (Supplementary Table S3). The main region containing NUMTs was on the largest scaffold (PGA_scaffold_1), where they are present as a tandem array encompassing a region of 640 Kb (Fig. 1F). Pseudogenisation was observed in some of the coding sequences in the mitochondrial assembly variants (Supplementary Fig. S3).

### Analysis of gene family expansion reveals proliferation of mobile elements

We found a total of 679 orthologous clusters with significant expansion or contractions (P < 0.05) across all the tick genomes tested, whether cell line- or whole tick-derived (Supplementary Fig. S4). Among these, 139 were expanded and 329 were contracted in the AVL/CTVM17 assembly. Notably, 53 (38%) of the expanded clusters contained PFAM annotations related to TEs such as transposases and reverse transcriptases. Among the ten largest expanded clusters (range of 19 to 178 copies), eight comprised TE-related genes (Supplementary Table S4), revealing a proliferation of mobile elements in the AVL/CTVM17 assembly. To determine if these expansions had occurred *in vitro*, we mapped the Illumina reads (Supplementary Methods, Table S1) from the *A. variegatum* cell lines (AVL/CTVM13 and AVL/CTVM17) and a subset of Cameroonian field-collected ticks (n = 5) to genes from the 53 expanded TE-related clusters, along with a random sample of 50 single-copy genes. Read coverage from the TE-related genes relative to single-copy genes was elevated across both cell lines and the ticks indicating that the TE expansions occurred prior to cell line development (Supplementary Table S5).

### TEs are major components of the AVL/CTVM17 genome

Combining *de novo* TE identification with automated and manual curation, we identified 958 distinct *bona fide* TE families across 28 superfamilies (Fig. 2A), revealing that TEs comprise at least 43.98% of the AVL/CTVM17 genome (3.78 Gb). The LINE order was the most prevalent (18.7%) (Fig. 2B), including members from distinct superfamilies (Fig. 2C). The LTR retrotransposons were the second most common category (11.3%), largely represented by the Gypsy superfamily, which was the most abundant in the genome and also the most diverse in the TE library (291 distinct families).

**Fig. 2.**
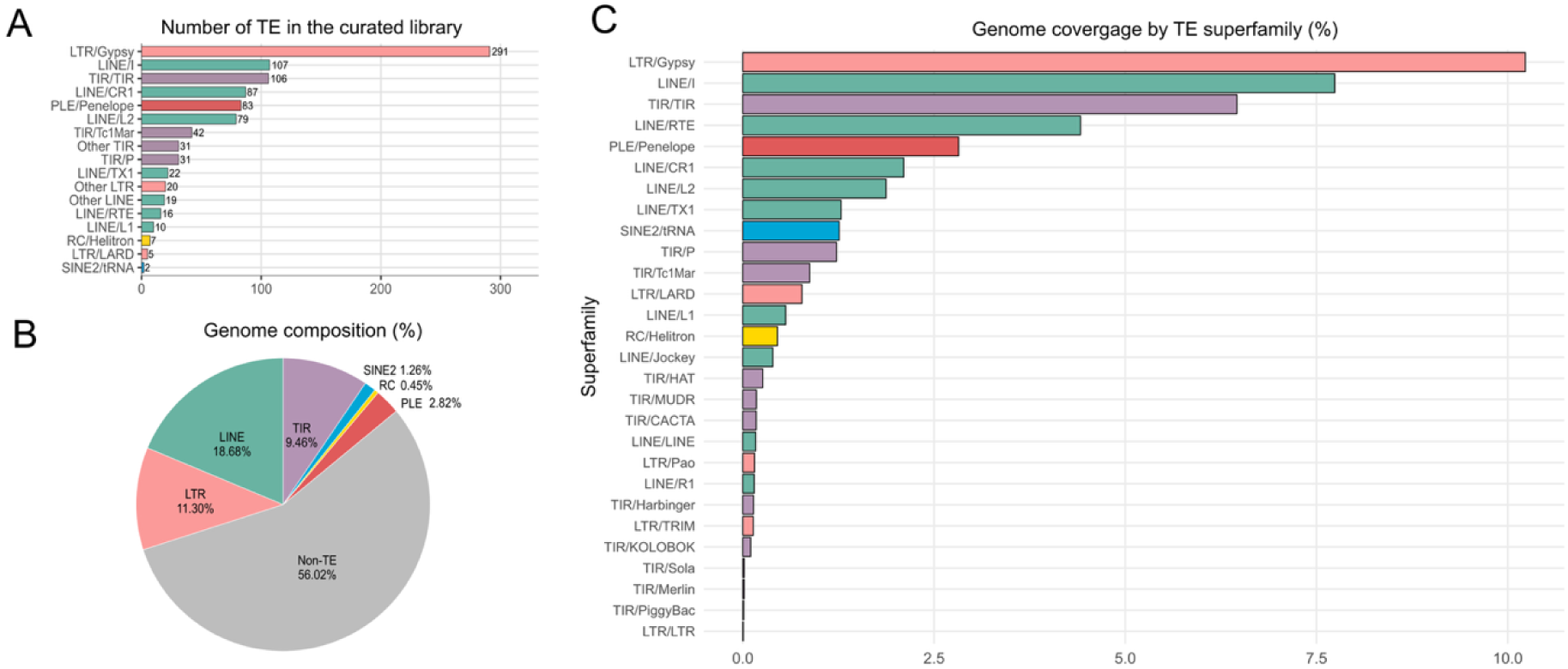
TEs constitute a large proportion of the *Amblyomma variegatum* genome. (A) Number of distinct families in each superfamily after curation of the TE library. (B) Genome proportions of the major TE orders. (C) Genome coverage of the major TE superfamilies. The TEs populating the *A. variegatum* genome include long interspersed nuclear element (LINE) retrotransposons, long terminal repeat (LTR) retrotransposons, terminal inverted repeat (TIR) transposons, Penelope-like (PLE) retrotransposons, short interspersed nuclear elements (SINEs) and rolling-circle (RC) transposons.

Remarkably, DNA transposons from the TIR order accounted for 9.5% of the genome. A substantial fraction (6.46%) corresponded to elements containing the terminal inverted repeats but lacking coding regions; these were designated here as TIR/TIR since they could not be assigned to any specific TIR superfamily. It is important to note that we did not screen for target-site duplications that would indicate whether these sequences are being replicated through transposition. These TIR-like elements were the most frequently found repetitive sequences (22%) in *I. ricinus*^27^.

Less common TEs were from the Penelope-like (2.82%), SINEs (1.26%) and rolling-circle (0.45%) orders. Additionally, we identified ten different palindromic sequence families ranging from 200 bp to 1.4 kb that covered 0.34% of the genome, although their mechanism of amplification is unclear.

### A near full-length *R. africae* chromosome sequence is integrated into the AVL/CTVM17 genome

Consistent with our previous observations^21^, Kraken2 classification of the PacBio assembly (the Hi-C assembly was too large for this analysis) revealed the presence of spotted fever group *Rickettsia* sequences. The absence of other bacterial sequences indicated a low likelihood of other bacterial LGT. We also performed Blobtools analysis on the smaller contigs (excluding the ten largest) from the Hi-C assembly, and did not identify any non-eukaryotic sequences, suggesting the absence of contaminating bacteria or DNA viruses.

By aligning the chromosome sequence of *R. africae* ESF-5 to the AVL/CTVM17 Hi-C assembly, we identified a region of approximately 4.5 Mb on the second largest scaffold (PGA_scaffold_2, the putative X chromosome) containing multiple insertions of *R. africae* sequence (Fig. 3A). Additional, minor (∼1 Kb) insertions were also detected on scaffolds 1, 6, 7 and 9, but the vast majority of *R. africae* sequences were concentrated on PGA_scaffold_2.

**Fig. 3.**
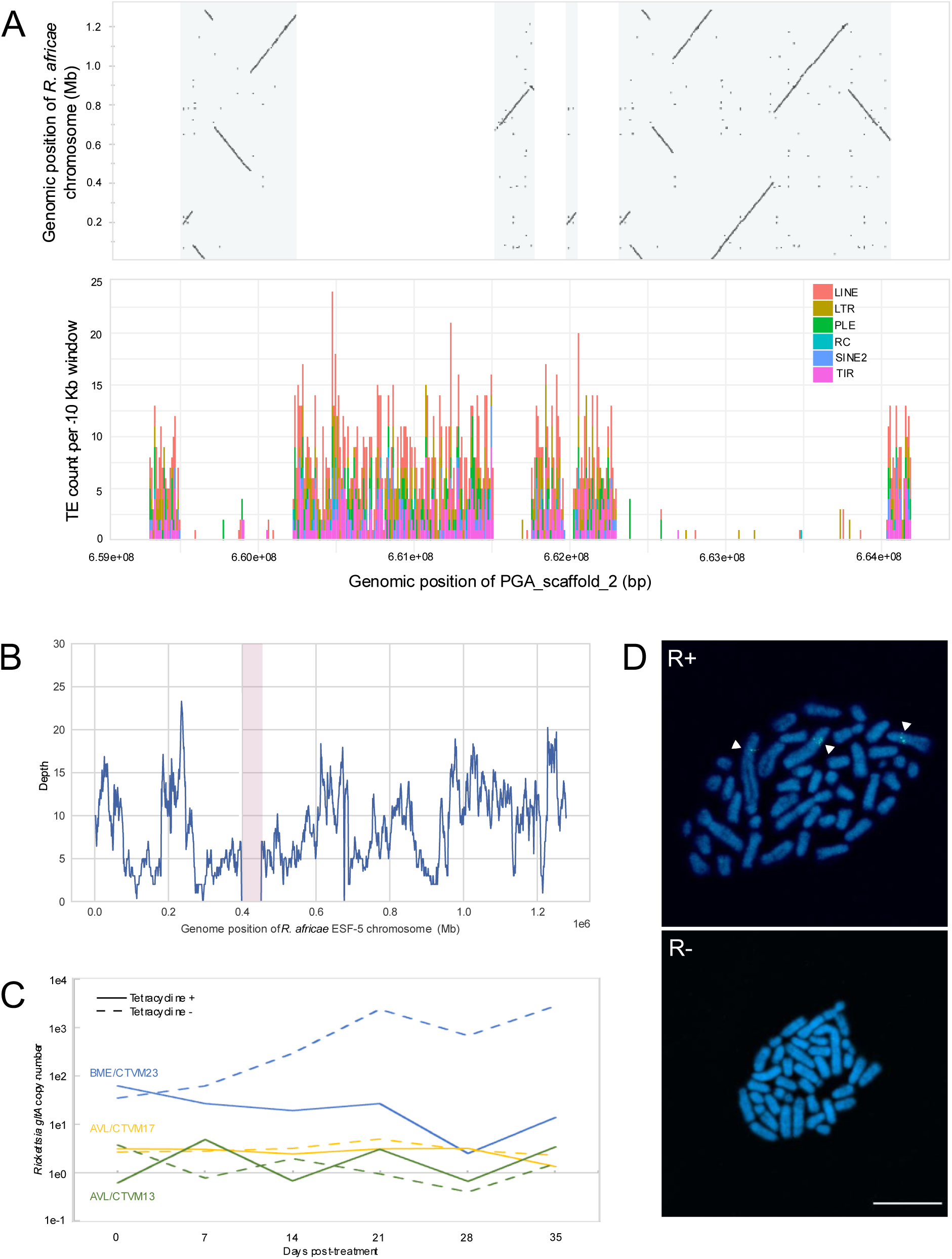
*Rickettsia africae* insertion sequence in the AVL/CTVM17 genome. (A) Alignment of the *R. africae* ESF-5 chromosome sequence to the second largest scaffold (PGA_scaffold_2) in the Hi-C assembly. The bottom panel depicts the counts of various TE groups across the indicated region. (B) Coverage plot of the PacBio sequence library across the *R. africae* ESF-5 chromosome sequence with a 1 Kb rolling window. The missing ∼54 Kb region is highlighted in pink. (C) *Rickettsia gltA* gene copy numbers in *A. variegatum* cells treated with tetracycline for 5 weeks or untreated. *Rhipicephalus microplus* (BME/CTVM23) cells infected with *Rickettsia raoultii* were used as controls to represent infection with live bacteria. (D) FISH on metaphase chromosome spreads from AVL/CTVM17 cells with probes targeting the entire length of the *R. africae* chromosome sequence (R+) -or only the missing 54 Kb region (R-). White triangles indicate chromosomes with hybridisation. The scale bar represents 20 µm.

We inspected the alignment of the AVL/CTVM17 PacBio and ONT sequence libraries on the Hi-C assembly and observed sequence reads spanning the junctions between the tick and bacterial insertion sequences (Supplementary Fig. S5), indicating that the presence of *R. africae* sequences within the Hi-C assembly is unlikely to be due to an assembly artefact.

We next determined the coverage of the PacBio, ONT and Illumina sequence libraries (Supplementary Methods, Table S1) produced from the AVL/CTVM17 and AVL/CTVM13 cell lines, as well as ticks from the Slovakia colony (n = 2 pools), across the *R. africae* ESF-5 genome. Reads from the sequence libraries mapped to nearly the entire length of the chromosome sequence of *R. africae* ESF-5, except for a region of approximately 54 Kb near the 400 Kb position (Fig. 3B and Supplementary Fig. S6). Thus, ∼95% of the *R. africae* chromosome is represented in the insertion sequence. However, coverage was notably uneven across the length of the *R. africae* chromosome, which is consistent with the extensive duplication and rearrangement of the insertion sequence in the Hi-C scaffold as seen in Fig. 3A, indicating that some regions were duplicated up to four times. Even though ∼60% of the scaffold comprises TEs, only a small number of TEs were present in the *R. africae* insertion (Fig. 3A, bottom panel). Similarly, TEs were not detected in the NUMT array (data not shown). Inspection of the BAM files from the alignments of the PacBio and ONT data from AVL/CTVM17 confirmed an absence of reads aligning to the *R. africae* chromosome at the 54-Kb missing region (Supplementary Fig. S7A). A closer examination of this gap revealed that several genes on the *R. africae* chromosome could be missing in the *R. africae* insertion sequence (Supplementary Fig. S7A). Accordingly, PCRs targeting two of these missing genes, *mutS* and *rpoH*, did not amplify the genes in the *A. variegatum* cell lines or Slovakian colony ticks (Supplementary Fig. S7B), confirming the absence of these sequences. We also did not find sequence reads that mapped to the pRa plasmid^28^ from *R. africae*, suggesting that pRa is not represented among the LGT, and consequently that live *R. africae* organisms are absent from the cell lines. This observation was further corroborated by an antibiotic treatment experiment, which did not affect the *Rickettsia gltA* copy number in either *A. variegatum* cell line over a period of 5 weeks’ continuous treatment (Fig. 3C). In contrast, tetracycline treatment reduced the *gltA* copy number in BME/CTVM23 tick cells infected with *R. raoultii,* representing a live rickettsial infection from the spotted fever group in which *R. africae* is classified.

### FISH reveals the *R. africae* sequence insertion on tick cell chromosomes

To visualise the insertion sequence on tick cell chromosomes, we designed a FISH experiment using probes targeting the full-length *R. africae* chromosome sequence (R+) or only the missing 54 Kb sequence (R-). The R+ probes clearly hybridised to the chromosomes in AVL/CTVM17 cells, whereas the R-probes did not (Fig. 3D). Hybridisation was occasionally observed on more than one chromosome, which could be explained by the high numbers of chromosomes present in most AVL/CTVM17 cells (Supplementary Fig S1) due to karyotype changes in long-term cultivated cell lines, resulting in variable chromosome numbers in each cell^25^. The distance between the hybridisation signals and the furthest end of the chromosomes, as a proportion of the length of the entire chromosome, was determined to be 0.67 (n = 20, standard deviation = 0.08, confidence interval = 0.03, alpha = 0.05), which is consistent with the approximate starting position of the main rickettsial insertion at 659.5 Mb (PGA_scaffold_2 length: 1099 Mb).

### Quantification of the pRa plasmid enables discrimination between *R. africae* bacteria and the insertion sequence in ticks

We next attempted to identify the presence of the *R. africae* insertion sequence in field-collected ticks and sought to discriminate it from sequences originating from active *R. africae* infections. Using a widely used *Rickettsia gltA* qPCR assay^29^, we assayed field-collected adult *A. variegatum* ticks collected from Angola (n = 2), Cameroon (n = 128), Mozambique (n = 6), Zambia (n = 3) and Zimbabwe (n = 2), and found that 95% were positive. However, the qPCR assay was unable to discriminate between rickettsial organisms or insertion sequence, since *gltA* is intact in the insertion sequence (see section for MLST genes below). Therefore, a subset of the positive ticks (n = 36) was selected for Illumina sequencing (Supplementary Methods, Table S1). These data, alongside Illumina reads from the *A. variegatum* cell lines, were mapped to the *A. variegatum* Hi-C scaffold containing the *R. africae* insertion (PGA_scaffold_2). The average sequencing depth was similar for all specimens (Fig. 4A, left). Restricting the mapping to the 4.5 Mb region containing the *R. africae* chromosomal insertion (i.e., 659 – 664 Mb position on PGA_scaffold_2) revealed high variability in the average sequencing depth across tick specimens.

**Fig. 4.**
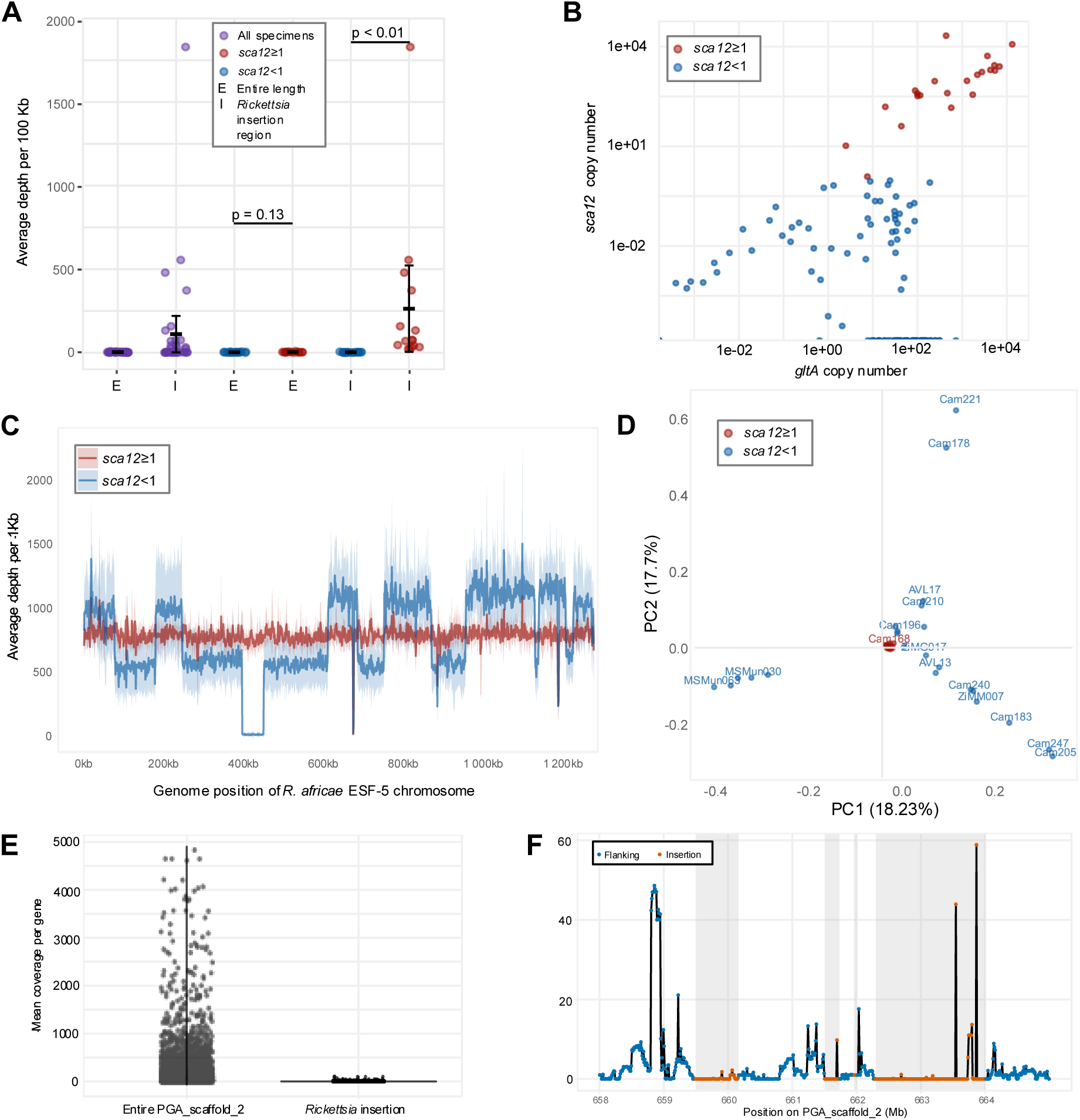
*Rickettsia africae* insertion and bacteria in *Amblyomma variegatum* ticks and cell line. (A) Left: Average sequence depth of sequence reads from *A. variegatum* ticks (n = 36) and cell lines (n = 2) aligned to the entire length of PGA_scaffold_2 from the AVL/CTVM17 Hi-C assembly or only to the region with the *R. africae* insertion (659.6 – 664.2 Mb). Right: Re-analysis of the average sequence depth was performed with specimens separated into groups based on a rickettsial plasmid gene (*sca12*) copy number: *sca12* ≥1 or *sca12* <1. Unpaired t-tests were performed for comparison of the means. Error bars represent mean ± 1 standard deviation (SD). (B) Scatter plot of *sca12* and *gltA* (a rickettsial chromosome gene) copy numbers assayed by qPCR from *A. variegatum* ticks (n = 141). (C) Coverage plot of tick sequence libraries mapped across *R. africae* ESF-5 chromosome sequence generated for a 1 Kb rolling window. The coverage was normalised to reads per million. Ribbons represent the mean coverage for *sca12* ≥1 (n = 15) or *sca12* <1 (ticks, n = 21; cell lines, n = 2) specimens ± 1 SD. (D) Principal component analysis (PCA) of SNP profiles generated from the mapped sequences. Key to sample labels are provided in Supplementary Methods Table S1). (E) Mean coverage per gene of ONT cDNA reads mapped to the entire length of PGA_scaffold_2 or just the *R. africae* insertion. (F) Coverage of ONT cDNA reads across the indicated region on PGA_scaffold_2.

We posited that the variability could be explained by the presence of a mix of ticks carrying only the *R. africae* insertion in the genome and those with active *R. africae* infections among the specimens. To test this hypothesis, we designed a qPCR assay targeting *sca12,* a gene unique to the *R. africae* pRa plasmid, which should only be present in the bacteria and not the insertion sequence. We were able to identify a population of ticks exhibiting one or more copies of *sca12* (i.e., *sca12* ≥1, Fig. 4B) per copy of a tick nuclear gene, *rpl6*, and a population in which *sca12* was undetectable or present in less than one copy (i.e., *sca12* <1) per copy of *rpl6*. Ticks with *sca12* ≥1 were also generally found to have higher *gltA* copy numbers. Based on this finding, we re-examined the average sequencing depth by grouping the specimens based on *sca12* ≥1 or *sca12* <1 (Fig. 4A, right). The average sequencing depth across the entire scaffold was similar for all specimens regardless of *sca12* copy numbers (P = 0.22). However, across the *R. africae* insertion region, ticks with *sca12* ≥1 displayed significantly higher average sequencing depth than ticks with *sca12* <1 (P = 0.048). The greater number of sequence reads aligning to the *R. africae* insertion region in the *sca12* ≥1 group supports the hypothesis that these ticks were infected with *R. africae* bacteria, or at minimum, contained *R. africae* DNA in the bloodmeal.

When aligned to the *R. africae* ESF-5 chromosome sequence, ticks with *sca12* ≥1 also exhibited even coverage across the entire length of the chromosome sequence (Fig. 4C), whereas ticks with *sca12* <1 displayed uneven coverage and a missing 54-Kb region (Fig. 4C). A principal component analysis (PCA) of SNP profiles revealed tight clustering of *sca12* ≥1 group specimens in contrast to the *sca12* <1 specimens (Fig. 4D), consistent with the lower genetic variability expected in *R. africae* organisms. Bacterial pangenome analysis of the assemblies generated from a subset of ticks (n = 8) and AVL/CTVM17 revealed the same set of missing genes (n = 72; 42 with COG functions, Supplementary Fig. S8, Table S6) shared between ticks with *sca12* <1 and the cell line, comprising the genes found in the 54-Kb missing region. Taken together, our data confirm that genetic differences exist between *R. africae* bacteria and the chromosomal insertion in the tick genome, which are likely to have arisen due to relaxed selection following the LGT. The pRa plasmid sequence including *sca12*, the missing 54-Kb region in the chromosomal insertion, and the distinct SNP profiles constitute discriminating features between *R. africae* infections and LGT.

### Some rickettsial ML ST genes in the *R. africae* insertion are disrupted

We performed a preliminary length-based screen to estimate the proportion of potential pseudogenised CDS. This analysis used CDS coverage as an initial indicator of sequence fragmentation and was not intended to generate a definitive catalogue of pseudogenes. Fragmented or truncated CDS were identified via BLASTn comparison with the CDS (n = 1,445) from the *R. africae* ESF-5 reference. In the AVL/CTVM17 *R. africae* insertion, only 646 CDS retained >80% coverage compared to the reference, suggesting a large number of CDS were potentially pseudogenised and non-functional in the insertion. A similar pattern was observed among Cameroon field-collected ticks from the pangenome analysis above, with an estimated 439.3 ± 108.8 potentially pseudogenised CDS in uninfected ticks (n = 3), compared with 8.2 ± 0.84 in infected ticks (n = 5). As the read coverages across the *R. africae* insertion region were generally lower in uninfected ticks (Fig. 4A), some fragmented CDS may have resulted from incomplete or mis-assemblies. Hence, we also performed full-length cDNA ONT sequencing for AVL/CTVM17, which showed minimal representation of transcripts across the *R. africae* insertion region on PGA_scaffold_2 (Fig. 4E&F), consistent with pseudogenisation. Multi-locus sequence typing (MLST) has been commonly used for classification of *Rickettsia* isolates30, and MLST genes have also been targeted for the development of rickettsial detection assays29, such as the *gltA* qPCR assay used for the screening of ticks in this study. We investigated the MLST genes in the *R. africae* genome insertion in the AVL/CTVM17 assembly for copy number variation and potential pseudogenisation. We identified one copy of intact *rrs*, two copies of intact *gltA*, and three copies each of fragmented or truncated *ompA*, *ompB* and *sca4*. Consequently, findings on the detection or classification of *Rickettsia* spp. in *A. variegatum* ticks based on these genes may be confounded by the presence of multiple copies and potential pseudogenisation, necessitating careful interpretation.

## Discussion

Here, to the best of our knowledge, we uncovered the first case of an almost-complete pathogen chromosome integrated into the genome of its arthropod vector. While confirming the presence of infectious particles, rather than the integration of viral elements into host genomes, is a well-recognised problem in virology^31,32^, this possibility is not routinely considered when screening vectors for other types of pathogens. Approximately two decades ago, the discovery of *Wolbachia*-derived LGT in multiple invertebrate species^12,33^ led to some concerns that PCR surveys of *Wolbachia* infections could lead to erroneous results, although it was assumed that LGT would consist of highly degraded pseudogenes and thus would be easily differentiable from active *Wolbachia* infections^17,18^. A general lack of recognised LGT from other bacterial symbionts in arthropod genomes, apart from capture of functional biosynthetic pathways in a few specific cases, appears to have led to a complacent outlook regarding the potential for LGT to complicate epidemiological investigations of vector-borne pathogens.

In contrast with most other terrestrial arthropod taxa, there is no evidence for widespread *Wolbachia* symbiosis in ticks, since most apparent infections result from the presence of insect parasitoids (*Ixodiphagus* spp.^34,35^) or filarial nematodes^36^. Instead, ticks are associated with a distinct spectrum of vertically-transmitted symbionts, including *Rickettsia* spp., *Coxiella*-like symbionts, *Francisella*-like symbionts, and *Candidatus* Midichloria spp^37,38^. In addition to the *R. africae* LGT identified here in *A. variegatum*, previous PCR screening of three tick cell lines derived from *Dermacentor* spp. amplified sequences from *Francisella*-like symbionts in the absence of microscopic evidence of bacterial infections^21^. This suggests that the situation in *A. variegatum* is not unique among ticks, further complicating epidemiological assessments of bacterial pathogens in these vectors, which is already clouded by the fact that tick microbiomes may contain both vertebrate pathogens and closely related arthropod-restricted symbionts with wide and overlapping distributions^39,40^.

Since ticks have the largest and most repetitive genomes among all arthropod vectors sequenced to date ^41^ ^42,43^, confirming the presence of LGT is especially challenging. Here, the use of tick cell lines was critical in the transition from a suspected case of LGT to its precise characterisation at the chromosomal level. In comparison with whole arthropods, visualisation of live bacteria in cell lines is more straightforward, as are antibiotic treatments to determine if molecular-based assays are potentially misleading. Cell lines also facilitate the production of relatively large amounts of axenic material for DNA isolations and are more conducive to the preparation of high molecular-weight DNA with low levels of contaminants, such as haem. This was essential to produce a high-quality assembly for the largest tick genome sequenced to date. However, a drawback of cell lines for genomic studies is the potential risk of genome rearrangements and duplications during serial passage^25^. While unlikely, it was possible that the LGT arose during maintenance of the tick colony used to generate the *A. variegatum* cell lines, or even during the development of primary cultures, and therefore had no epidemiological relevance.

This possibility was excluded by Illumina sequencing of ticks from a different colony to the one used for cell line generation, as well as short-read sequencing of over 30 wild-caught ticks from across sub-Saharan Africa, demonstrating that the LGT is a widespread, probably universal, feature of the genomes of *A. variegatum* ticks. The AVL/CTVM17 genome appeared to have a very similar structure to the genome of the source tick, and both the cell line genome and the assembly from the colony-derived ticks were resolved into 10 pseudochromosomes (i.e., one chromosome fewer than the expected haploid karyotype^24^). Future studies could incorporate optical mapping to determine if a mis-assembly of two small chromosomes has occurred. Irrespective of this discrepancy, the AVL/CTVM17 cell line is a rich resource for facilitating research on *A. variegatum* – one of the most important ticks affecting domestic ruminants in sub-Saharan Africa and the Caribbean, and the primary vector not only of *R. africae* but also of *Ehrlichia ruminantium* (agent of heartwater disease in ruminants^44,45^) and Dugbe virus (agent of a rare but highly virulent zoonosis)^46^.

While the TE content of the AVL/CTVM17 genome represented almost half of the assembly, this was substantially lower than in the castor bean tick, *Ixodes ricinus*, in which it reaches 69%^43^. The large array of NUMTs exhibiting relatively low levels of degradation compared to the *A. variegatum* mitogenome were a distinctive feature of the AVL/CTVM17 genome that has not been reported for other ticks, although large NUMTs are present in the genome of another chelicerate, *Limulus polyphemus*^47^. Similarly to the *R. africae* insertion, the NUMTs of *A. variegatum* have the potential to cause errors in molecular epidemiological and taxonomic studies^48^, since tick mitochondrial loci rather than nuclear genes are frequently targeted in these investigations^49^.

The *R. africae* insertion on the putative X chromosome of *A. variegatum* is reminiscent of the extensive *Wolbachia* LGT on the X chromosome of the adzuki bean beetle, *Callosobruchus chinensis*, which was estimated to represent ∼30% of the *Wolbachia* genome and exists in parallel with live *Wolbachia* infections in this species^50^. As with the *R. africae* insertion, transcriptional activity of this *Wolbachia* LGT was minimal. However, it appears to be more extensively degraded than the *R. africae* insertion, and LGT content exhibits high variability between beetle populations. Notably, in animals with XY or XO sex determination systems, the X chromosome can only undergo recombination in females^51^. This may facilitate retention of nonfunctional LGT following the original integration event, which in the case of *A. variegatum*, may have been relatively recent considering the comparatively limited pseudogenisation observed across the insertion and relatively low accumulation of TEs within the inserted region.

In conclusion, our findings are crucial in broadening awareness of prokaryote-to-eukaryote LGT of Rickettsiales origin from the ubiquitous symbiont *Wolbachia*, of which clinicians and epidemiologists outside the arbovirus and filariasis fields are generally unaware, to a pathogenic species of *Rickettsia*, with important implications for PCR-based pathogen screening in vectors. As microbial “contaminants” are often removed during genome assembly, and chromosomal-scale genomes have only recently become available for arthropods, further cases of LGT involving pathogenic bacteria may await discovery. In the meantime, pathogen detection by PCR alone in any vector lacking a high-quality genomic reference that has been scanned for LGT could generate invalid conclusions. Fortunately for epidemiological studies of ATBF, pRa appears to be a convenient target for low-cost molecular screening of genuine *R. africae* infections in ticks. The potential for epidemiological misconceptions is highlighted by the original *R. africae* genome study, in which plasmid-free strains were reported from some populations of *A. variegatum* based on PCR screening, despite the universal presence of pRa in clinical isolates^28^. While the existence of plasmid-free strains is theoretically possible, it is much more probable that these ticks harboured LGT but lacked current *R. africae* infections. Nevertheless, genomic surveys of *A. variegatum* and other vectors of *R. africae* across sub-Saharan Africa and the Caribbean are warranted to understand variation in LGT, and potentially pRa prevalence, before the epidemiology of ATBF and the biology of *R. africae* can be fully understood.

## Methods

Full, detailed methods and information for all materials sequenced are provided in the Supplementary Methods.

### Ticks, tick cell lines and *Rickettsia* spp

*Amblyomma variegatum* ticks were obtained from a second laboratory generation maintained at the Institute of Zoology and Center of Biomedical Sciences of the Slovak Academy of Sciences, Bratislava, Slovakia. This was derived from a laboratory colony at the French Agricultural Research Centre for International Development (CIRAD), UMR ASTRE, F-97170 Petit-Bourg, Guadeloupe, France. Field-derived *A. variegatum* ticks were also collected from cattle during routine de-ticking for welfare reasons by veterinarians in Ngaoundéré, Adamawa Region of Cameroon, during a previous project^52^. Finally, DNA extracts from ticks collected from southern Africa for a previous study^22^ were also used here.

The two *A. variegatum* cell lines AVL/CTVM13^53^ and AVL/CTVM17^54^ were maintained at 32°C in L-15/L-15B and L-15/H-Lac/L-15B medium, respectively^54^; whereas the *Rhipicephalus microplus* cell line BME/CTVM23 was maintained in L-15 medium at 32 °C as described previously^55^. The *Rickettsia raoultii* strain Białystok1 was cultured in BME/CTVM23 cells as previously reported^21,56^. Positive control *R. africae* DNA was kindly provided by Prof. Marcelo Labruna, University of Sao Paulo, Brazil.

### Antibiotic treatment assay

Duplicate cultures of AVL/CTVM13, AVL/CTVM17 and *R. raoultii*-infected BME/CTVM23 cells, incubated at 32°C with a weekly medium change, were either treated weekly with 100 µg/ml tetracycline hydrochloride (Sigma) or left untreated as described in the Supplementary Methods. DNA was extracted from sampled cells prior to treatment and weekly thereafter up to day 35 and subjected to quantitative PCR assays targeting fragments of the *Rickettsia* citrate synthase gene^57^ (*gltA*) and the tick *rpl6* gene^58^. The counts obtained for the *gltA* gene were normalised to the tick *rpl6* gene.

### Fluorescence *in situ* hybridisation (FISH)

Biotin-labelled myTags Custom probes were designed and produced by Daicel Arbor Biosciences, US. Briefly, metaphase chromosome spreads were prepared from a 3-day-old culture of AVL/CTVM17 cells following colcemid treatment as described in Supplementary Methods. The chromosome spreads were denatured at 70°C and incubated with the FISH probes overnight at 37°C for hybridisation. Post-hybridisation, the chromosome spreads were labelled with Alexa Fluor™ 488 streptavidin conjugate (Invitrogen, UK). The chromosome spreads were visualised and imaged at the Liverpool Centre for Cell Imaging (CCI), University of Liverpool, using a Zeiss Axio Observer.Z1/7 with LSM900 confocal microscope.

### Genomic sequencing, assembly and annotation

For PacBio sequencing, DNA was extracted from a single AVL/CTVM17 culture (p125). PacBio library preparation and sequencing were performed in the Centre for Genomic Research at the University of Liverpool (CGR). Assembly of the mitochondrial genome from the PacBio dataset has been previously described^26^. Oxford Nanopore Technologies (ONT)-based sequencing was performed twice for AVL/CTVM17 cells (p125-127). The first run used the DNA preparation as above for library preparation according to SQK-LSK109 protocol. The second run was performed at the University of Nottingham using a previously published ultra-long DNA protocol^59^. Data from both ONT runs were combined for assembly. For Illumina sequencing, DNA was extracted from AVL/CTVM13 (p155) and AVL/CTVM17 (p139) cells, frozen salivary glands harvested from ten each of female and male Slovakia colony ticks (pool 7D and 9D) or from half of individual field-collected Cameroon ticks (male and female). DNA extracts from ticks collected from Southern Africa for a previous study^22^ was also used for Illumina sequencing. Illumina library preparation and sequencing was performed at the CGR. Hi-C sequencing was performed using tissues from three male ticks from the Slovakia colony and cells from a single AVL/CTVM17 culture (p139). DNA in the material was cross-linked with 1% formaldehyde and delivered to Phase Genomics (Seattle, WA, USA) for DNA extraction, Proximo^TM^ Hi-C library preparation and Illumina S4 Novaseq sequencing. Genome assembly, quality control and annotation methods were given in the Supplementary Methods.

### RNA extraction and cDNA sequencing

Whole cell RNA was extracted from single AVL/CTVM17 culture (p139) using the RNAeasy Mini Kit (Qiagen, UK). The NEBNext® Single Cell/Low Input cDNA Synthesis & Amplification Module (New England Biolabs, UK) was used for generating high-quality, full-length cDNA, which was used for library preparation according to the SQK-NBD114-24 protocol and sequenced with FLO-MIN114 (R 10.4.1) flow cells for GridION (ONT, UK). Raw POD5 files were base-called with ONT Dorado v7.2.13 using the high-accuracy, 400 bps model.

### Phylogenetic tree inference

The detailed methods for phylogenetic inference and the accession numbers for all the tick genome assemblies used in the analysis are given in the Supplementary Methods. Briefly, single-copy orthologue genes were obtained from each tick genome and aligned. The concatenated alignments were then used for phylogenetic tree building.

### Gene family expansion analysis

Orthologous clusters from the longest isoform of each gene for each tick species were identified and provided to CAFE5^60^ for gene family contraction/expansion analysis. Gene families related to TEs were identified from the Pfam annotations. To test if the expansion of TE clusters was unique to cell lines, the ratio of average Illumina read coverage per base for the identified TE genes to 50 randomly-selected single-copy genes was determined for the cell lines and representative ticks as described in the Supplementary Methods.

### Annotation of TEs

RepeatModeler v2.0.3^61^ was employed to predict *de novo* TE in the *A. variegatum* assembly using the LTRstruct option. Subsequently, the library was curated using automated tools and manual inspection as described in the Supplementary Methods. The classification of TE families was either obtained using the RepeatClassifier module^61^ or by a visual inspection of structure and protein domains. The curated TE library was used in RepeatMasker v4.1.1^62^ to get the complete TE annotation. For TE coverage estimates, we excluded PALINDROME annotations and retained only high-confidence TE annotations ≥50 bp.

### *Rickettsia africae* and nuclear mitochondrial DNA (NUMT) insertion sequence analysis

The *R*. *africae* ESF-5 reference assembly, *A. variegatum* mitochondrion assembly (Accession: PZ344169.1)^26^, and Illumina short reads from ticks and tick cell lines were aligned to the AVL/CTVM17 cell line Hi-C scaffolds. Mapping coverage, SNPs calling and PCA were performed as described in the Supplementary Methods. Unpaired t-tests were performed for comparison of the means, assuming a normal distribution as determined from the Shapiro-Wilk normality test.

### *Rickettsia africae* genome assembly and pangenome analysis

Sequencing reads from the AVL/CTVM17 cell line, field-collected ticks and *Amblyomma sparsum* AM14 (Genbank SRA: SRR24067184)^63^ were mapped to the *R. africae* ESF-5 reference and then assembled using the appropriate sequence assemblers for pangenomic analysis as described in the Supplementary Methods.

### Identifying potential pseudogenes

To identify potential pseudogenes, the CDS predicted from the *R. africae* insertion region in the AVL/CTVM17 Hi-C assembly and also the pangenome workflow from the field-collected ticks were compared to the CDS from the *R. africae* ESF-5 reference assembly using BLASTn. Those CDS with >95% percentage identity to the *R. africae* ESF-5 reference CDS were classified as near full-length (>80% coverage of the reference CDS), whereas those that exhibited <80% coverage of the reference CDS were defined as truncated/fragmented (*i.e*., potential pseudogenes). The proportion of truncated/fragmented CDS was determined for field-collected ticks previously classified as infected (*sca12* ≥1) or uninfected (*sca12* <1). The MLST gene copy number and fragmentation were identified from the AVL/CTVM17 Hi-C assembly.

## Acknowledgements

This research was funded by the United Kingdom Biotechnology and Biological Sciences Research Council (grant no. BB/P024270/1) and the Wellcome Trust (grant no. 223743/Z/21/Z). AB was funded by the European Union’s Horizon 2020 research and innovation programme under the Marie Skłodowska Curie grant agreement No. H2020 MSCA ITN 2015 675752, and AMA was funded by the Republic of Iraq Ministry of Education & Scientific Research (doctoral scholarship).

All tick cell lines used in the study were sourced from the Tick Cell Biobank at the University of Liverpool. We acknowledge the assistance of Thomas Waring from the Centre for Cell Imaging (CCI), University of Liverpool, for acquiring the confocal microscopy images; Grace Ward, University of Liverpool, for performing the qPCR analysis; and Glen Robinson, University of Liverpool, for conducting the tetracycline treatment experiment. We are grateful to Andrea Betancourt (University of Liverpool) for helpful comments on the manuscript.

## Data Availability

All sequencing data generated from this study has been deposited in the European Nucleotide Archive (Project accession: PRJEB76275).

## Supplementary information

Full methods are provided in the Supplementary Methods.

Supplementary figures and tables are available in the Supplementary File.

---

**Table S1.** Sequence libraries and assemblies generated from *A. variegatum* AVL/CTVM17 cell line and ticks.

|  | PacBio | ONT 1D | ONT Ultra-long | Illumina | Hi-C |
| --- | --- | --- | --- | --- | --- |
| **AVL/CTVM17** |  |  |  |  |  |
| Num of reads | 6,467,236 | 4,009,365 | 7,388,316 | 515,302,182 | #2,437,312,053 |
| Total bases (Gb) | 92.45 | 19.73 | 41.12 | ~77 | 731.19 |
| N50 (Gb) | 14,283 | 15,005 | 96,874 | - | *2,517,172 |
| **Slovakia ticks male AV123** |  |  |  |  |  |
| Num of reads | - | - | - | - | #411,374 |
| Total bases (Gb) | - | - | - | - | 0.0617 |
| N50 (Gb) | - | - | - | - | *874,674 |

**Sequence assemblies from AVL/CTVM17**
| Assemblies | Number of contigs | Total size (Gb) | N50 | GC (%) | BUSCO (%) |
| --- | --- | --- | --- | --- | --- |
| PacBio | 21,134 | 8.35 | 874,674 bp | 47.28 | 94 |
| ONT (ONT 1D + ONT Ultra-Long) | 60,412 | 5.12 | 386,808 bp | 47.39 | 84.7 |
| Hi-C scaffolded (PacBio + Hi-C) | 721 | 8.61 | 1.02 Gb | 47.28 | 93.3 |

**Fig. S1.**
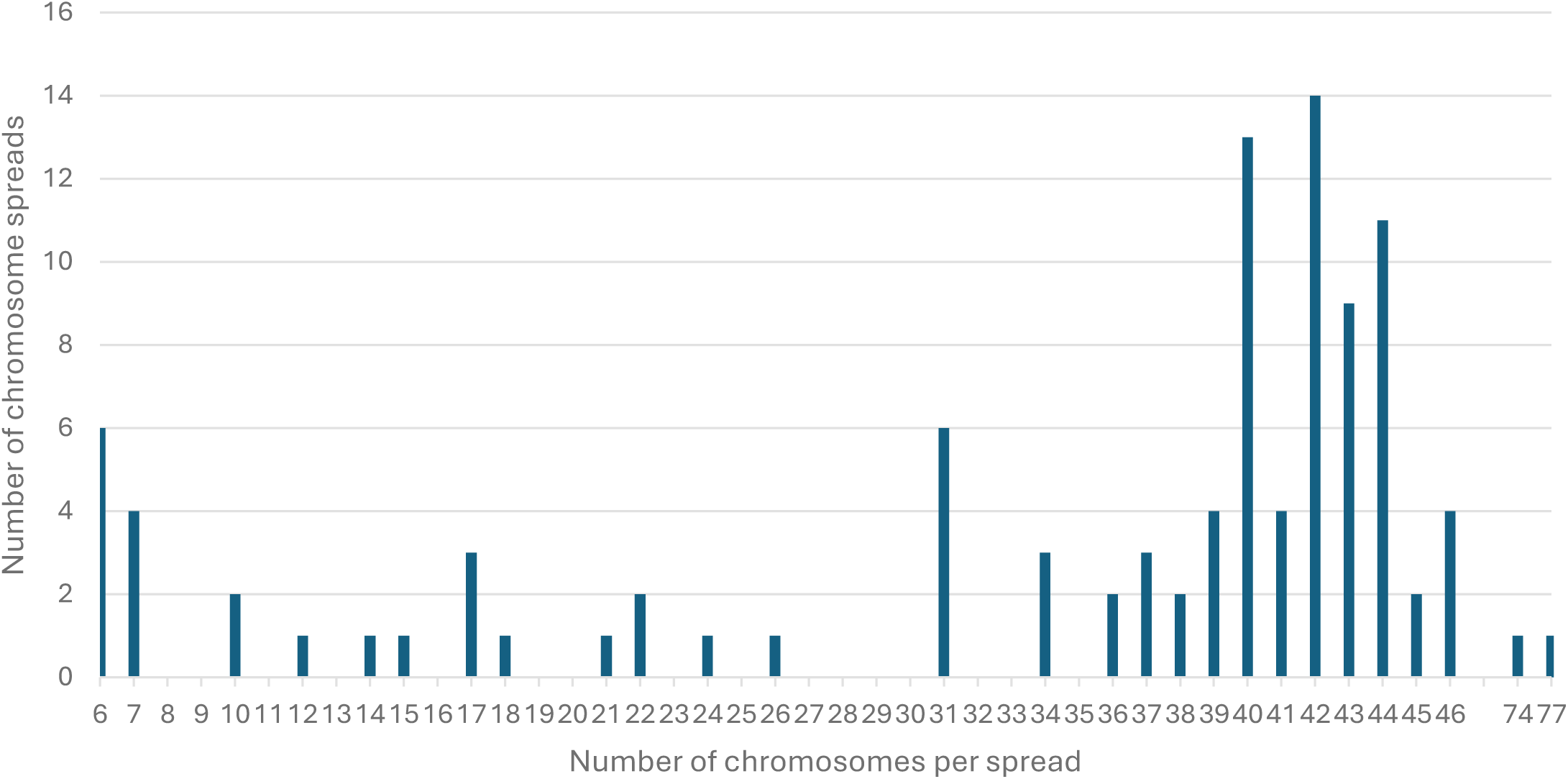
Chromosome numbers in AVL/CTVM17 metaphase chromosome spreads.

**Fig. S2.**
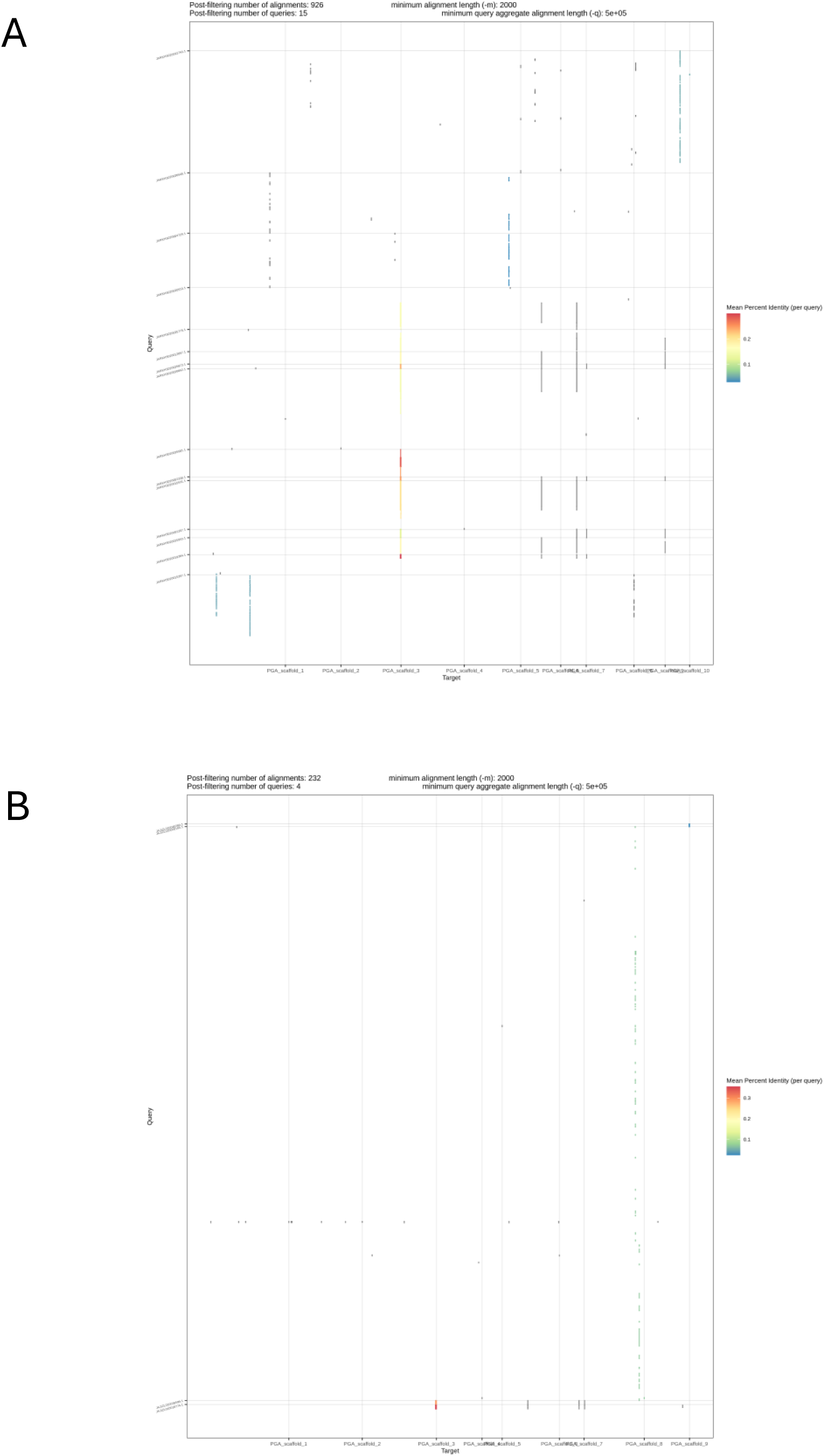
Alignment of genome assemblies from (A) Amblyomma americanum (ASM3014330v2) and (B) Amblyomma maculatum (ASM2396939v1) to AVL/CTVM17 assembly showed limited identity and synteny.

**Table S2.** Collinearity analysis of ten largest scaffolds in *A. variegatum* Hi-C assembly.

| Gene block | Score | E-value | Num of genes | Scaffolds | Orientation | Size in block 1 (bp) | Proportion in scaffold (%) | Size in block 2 (bp) | Proportion in scaffold (%) |
| --- | --- | --- | --- | --- | --- | --- | --- | --- | --- |
| 0 | 288 | 2.1E-12 | 6 | PGA_scaffold_1&PGA_scaffold_1 | minus | 72007 | 0.01 | 823389 | 0.06 |
| 1 | 231 | 7.2E-08 | 5 | PGA_scaffold_1&PGA_scaffold_1 | minus | 445795 | 0.03 | 1545446 | 0.12 |
| 2 | 262 | 1.3E-08 | 6 | PGA_scaffold_1&PGA_scaffold_9 | plus | 31130 | 0.00 | 653000 | 0.13 |
| 3 | 236 | 6.4E-10 | 6 | PGA_scaffold_1&PGA_scaffold_9 | plus | 31130 | 0.00 | 1106459 | 0.22 |
| 4 | 233 | 6.3E-08 | 6 | PGA_scaffold_1&PGA_scaffold_9 | plus | 31130 | 0.00 | 1091510 | 0.22 |
| 5 | 215 | 5.2E-07 | 5 | PGA_scaffold_1&PGA_scaffold_9 | plus | 31130 | 0.00 | 640558 | 0.13 |
| 6 | 318 | 5.4E-13 | 7 | PGA_scaffold_2&PGA_scaffold_2 | plus | 1774880 | 0.16 | 1018571 | 0.09 |
| 7 | 259 | 8.5E-10 | 6 | PGA_scaffold_2&PGA_scaffold_2 | plus | 1529762 | 0.14 | 1674661 | 0.15 |
| 8 | 366 | 0.0E+00 | 9 | PGA_scaffold_3&PGA_scaffold_3 | plus | 962783 | 0.09 | 1150966 | 0.11 |
| 9 | 349 | 3.9E-20 | 9 | PGA_scaffold_3&PGA_scaffold_3 | plus | 962783 | 0.09 | 1876872 | 0.18 |
| 10 | 318 | 5.2E-16 | 8 | PGA_scaffold_3&PGA_scaffold_3 | plus | 1117540 | 0.11 | 1347244 | 0.13 |
| 11 | 296 | 1.1E-12 | 7 | PGA_scaffold_3&PGA_scaffold_3 | plus | 916523 | 0.09 | 353335 | 0.03 |
| 12 | 287 | 3.3E-12 | 7 | PGA_scaffold_3&PGA_scaffold_3 | plus | 918564 | 0.09 | 1003532 | 0.10 |
| 13 | 265 | 3.0E-17 | 8 | PGA_scaffold_3&PGA_scaffold_3 | plus | 674814 | 0.06 | 2389557 | 0.23 |
| 14 | 254 | 4.2E-12 | 7 | PGA_scaffold_3&PGA_scaffold_3 | plus | 674814 | 0.06 | 1351561 | 0.13 |
| 15 | 253 | 7.5E-11 | 7 | PGA_scaffold_3&PGA_scaffold_3 | plus | 259210 | 0.02 | 1862441 | 0.18 |
| 16 | 235 | 7.7E-09 | 6 | PGA_scaffold_3&PGA_scaffold_3 | plus | 715166 | 0.07 | 1219692 | 0.12 |
| 17 | 214 | 2.3E-08 | 5 | PGA_scaffold_3&PGA_scaffold_3 | plus | 625016 | 0.06 | 500281 | 0.05 |
| 18 | 223 | 7.0E-09 | 6 | PGA_scaffold_3&PGA_scaffold_3 | minus | 259210 | 0.02 | 1297397 | 0.12 |
| 19 | 216 | 4.8E-10 | 6 | PGA_scaffold_3&PGA_scaffold_3 | minus | 259210 | 0.02 | 1518507 | 0.15 |
| 20 | 206 | 8.0E-06 | 5 | PGA_scaffold_3&PGA_scaffold_3 | minus | 218858 | 0.02 | 871867 | 0.08 |
| 21 | 203 | 2.3E-06 | 5 | PGA_scaffold_3&PGA_scaffold_3 | minus | 218858 | 0.02 | 893225 | 0.09 |
| 22 | 226 | 0.0E+00 | 5 | PGA_scaffold_3&PGA_scaffold_7 | minus | 218858 | 0.02 | 234104 | 0.03 |
| 23 | 350 | 5.7E-20 | 10 | PGA_scaffold_9&PGA_scaffold_9 | plus | 560862 | 0.11 | 714714 | 0.14 |
| 24 | 217 | 3.4E-07 | 5 | PGA_scaffold_9&PGA_scaffold_9 | minus | 392608 | 0.08 | 600425 | 0.12 |

**Fig. S3.**
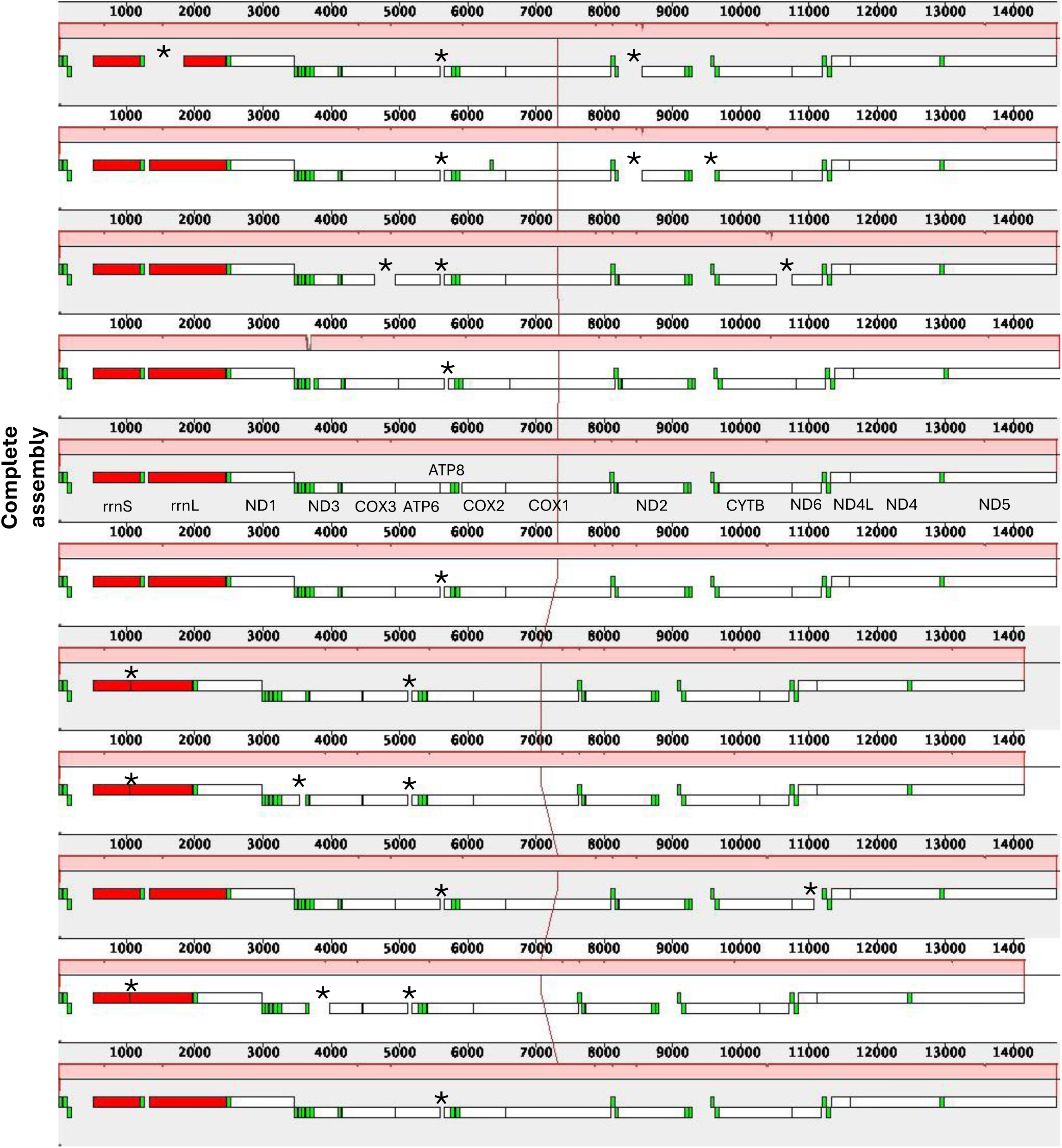
ProgressiveMauve alignment of complete mitochondrial assembly and multiple variants from *A. variegatum* cell line, AVL/CTVM17. Assemblies were produced from the MitoHifi pipeline using PacBio HiFi reads. Protein coding genes (white blocks) were labelled for the complete assembly. Ribosomal RNA genes were coloured in red. Transfer-RNA genes were coloured in green. Coding sequences with disruptions were indicated with asterisks.

**Table S3.** Starting and ending positions of NUMTs on the Hi-C scaffolds.

| Scaffold | Starting position (bp) | Ending position (bp) | Size (bp) |
| --- | --- | --- | --- |
| PGA_scaffold_1 | 667581615 | 668220612 | 638997 |
| PGA_scaffold_1 | 853304401 | 853306062 | 1661 |
| PGA_scaffold_1 | 853305303 | 853305867 | 564 |
| PGA_scaffold_1 | 853306063 | 853313876 | 7813 |
| PGA_scaffold_1 | 853310744 | 853311360 | 616 |
| PGA_scaffold_1 | 893771853 | 893772270 | 417 |
| PGA_scaffold_1 | 926018320 | 926018494 | 174 |
| PGA_scaffold_2 | 0 | 3850 | 3850 |
| PGA_scaffold_2 | 3223 | 3653 | 430 |
| PGA_scaffold_2 | 3850 | 14634 | 10784 |
| PGA_scaffold_2 | 8749 | 9190 | 441 |
| PGA_scaffold_2 | 368500004 | 368503133 | 3129 |
| PGA_scaffold_2 | 368860747 | 368863873 | 3126 |
| PGA_scaffold_2 | 494371025 | 494371998 | 973 |
| PGA_scaffold_3 | 187269661 | 187270045 | 384 |
| PGA_scaffold_3 | 904779720 | 904779958 | 238 |
| PGA_scaffold_3 | 904779945 | 904780270 | 325 |
| PGA_scaffold_4 | 269735357 | 269752903 | 17546 |
| PGA_scaffold_4 | 273327440 | 273339336 | 11896 |
| PGA_scaffold_4 | 276371406 | 276374079 | 2673 |
| PGA_scaffold_4 | 288287668 | 288312578 | 24910 |
| PGA_scaffold_4 | 362698855 | 362699829 | 974 |
| PGA_scaffold_4 | 365857221 | 365858194 | 973 |
| PGA_scaffold_5 | 572792293 | 572799146 | 6853 |
| PGA_scaffold_5 | 576111133 | 576112310 | 1177 |
| PGA_scaffold_6 | 279453030 | 279526519 | 73489 |
| PGA_scaffold_7 | 339113892 | 339114290 | 398 |
| PGA_scaffold_7 | 339114958 | 339115188 | 230 |
| PGA_scaffold_7 | 590122674 | 590122886 | 212 |
| PGA_scaffold_7 | 700374363 | 700374536 | 173 |
| PGA_scaffold_7 | 700374363 | 700374565 | 202 |
| PGA_scaffold_7 | 702425490 | 702425663 | 173 |
| PGA_scaffold_7 | 702425490 | 702425692 | 202 |
| PGA_scaffold_8 | 349692940 | 349701905 | 8965 |
| PGA_scaffold_8 | 349700138 | 349700747 | 609 |
| PGA_scaffold_8 | 349707074 | 349708279 | 1205 |
| PGA_scaffold_8 | 351670603 | 351671432 | 829 |
| PGA_scaffold_8 | 422114580 | 422115810 | 1230 |
| PGA_scaffold_8 | 428771961 | 428772672 | 711 |
| PGA_scaffold_8 | 472399526 | 472399890 | 364 |
| PGA_scaffold_8 | 472400807 | 472401353 | 546 |

**Fig. S4.**
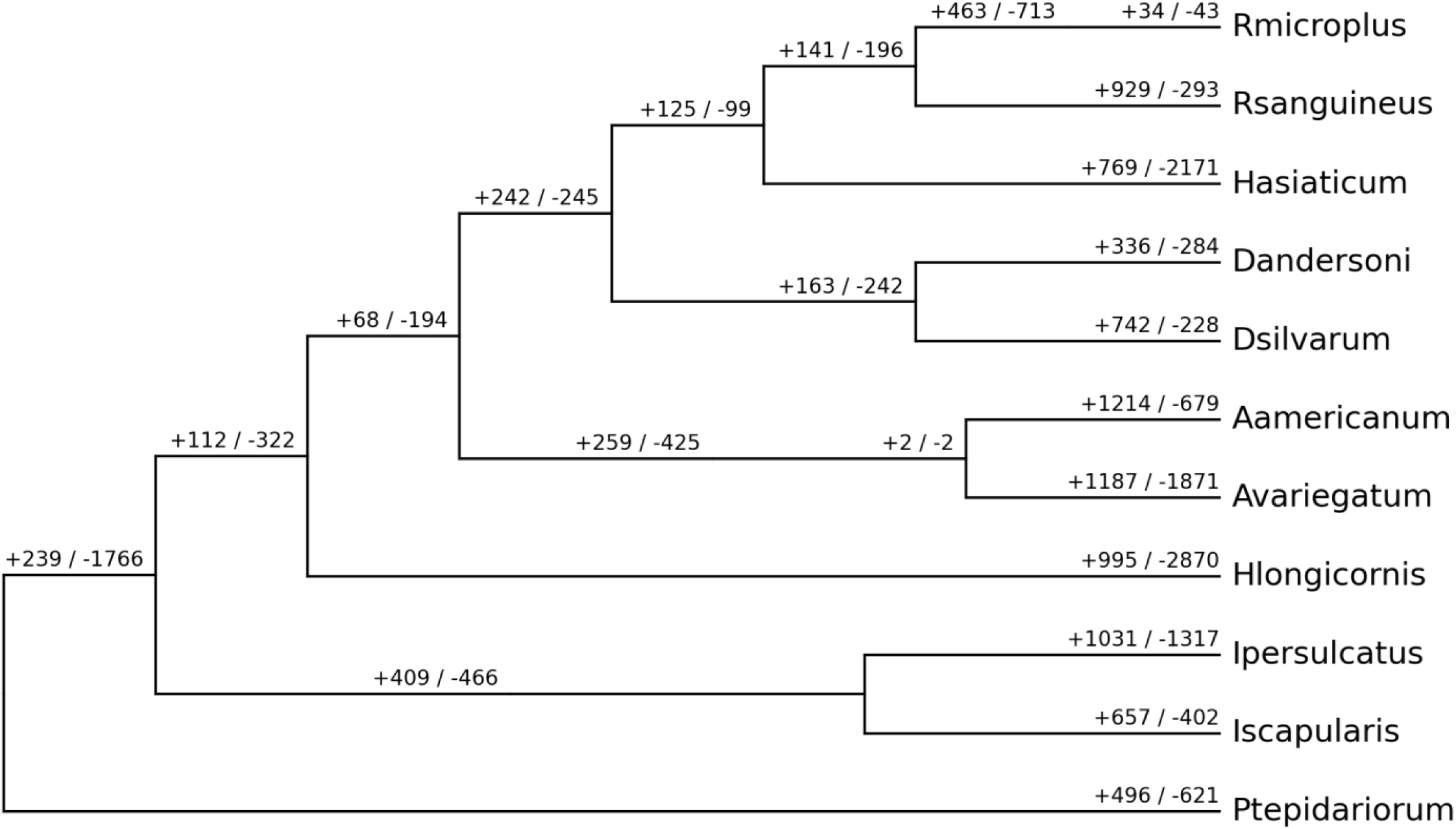
Gene families/orthologous clusters with expansion or contraction regardless of significance.

**Table S4.** Top ten expanded gene families/orthologous clusters in *A. variegatum* genome assembly (P<0.05).

| Orthogroup | Copies | EM_PFAM | EM_desc |
| --- | --- | --- | --- |
| OG0000016 | 178 | DUF4371,Dimer_Tnp_hAT | None |
| OG0000069 | 121 | Exo_endo_phos_2,RNase_H,RV<br>T_1 | Endonuclease-reverse<br>transcriptase |
| OG0000003 | 81 | Tnp_P_element | Transposase protein |
| OG0000029 | 60 | RVT_1,rve | transposition, RNA-mediated |
| OG0000254 | 39 | Not assigned | Not assigned |
| OG0000017 | 37 | DYW_deaminase,PPR,PPR_2 | oxidoreductase activity, oxidizing<br>metal ions with flavin as acceptor |
| OG0000038 | 31 | RVT_1,rve | transposition, RNA-mediated |
| OG0000374 | 26 | CENP-B_N,DDE_1,HTH_Tnp_Tc5 | Tigger transposable element<br>derived 6 |
| OG0000032 | 22 | Tnp_P_element | None |
| OG0000707 | 19 | THAP,Tnp_P_element | Transposase protein |

**Table S5.**
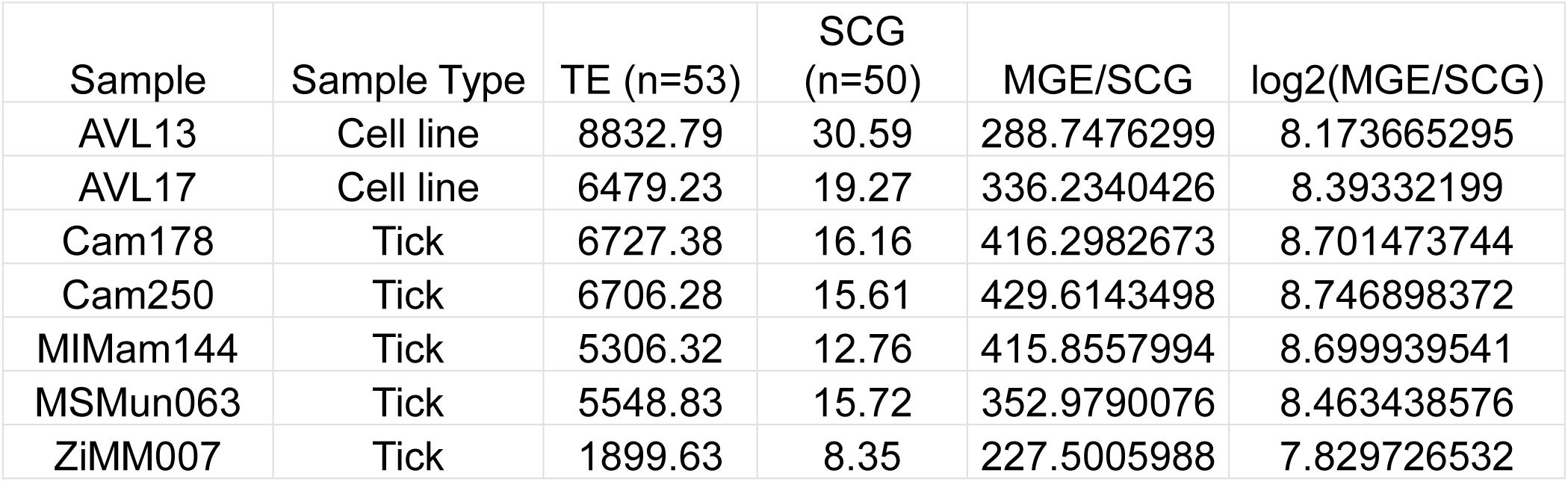
Average coverage per base for TE genes and single copy genes (SCG)

| Sample | Sample Type | TE (n=53) | SCG<br>(n=50) | MGE/SCG | log2(MGE/SCG) |
| --- | --- | --- | --- | --- | --- |
| AVL13 | Cell line | 8832.79 | 30.59 | 288.7476299 | 8.173665295 |
| AVL17 | Cell line | 6479.23 | 19.27 | 336.2340426 | 8.39332199 |
| Cam178 | Tick | 6727.38 | 16.16 | 416.2982673 | 8.701473744 |
| Cam250 | Tick | 6706.28 | 15.61 | 429.6143498 | 8.746898372 |
| MIMam144 | Tick | 5306.32 | 12.76 | 415.8557994 | 8.699939541 |
| MSMun063 | Tick | 5548.83 | 15.72 | 352.9790076 | 8.463438576 |
| ZiMM007 | Tick | 1899.63 | 8.35 | 227.5005988 | 7.829726532 |

**Fig. S5.**
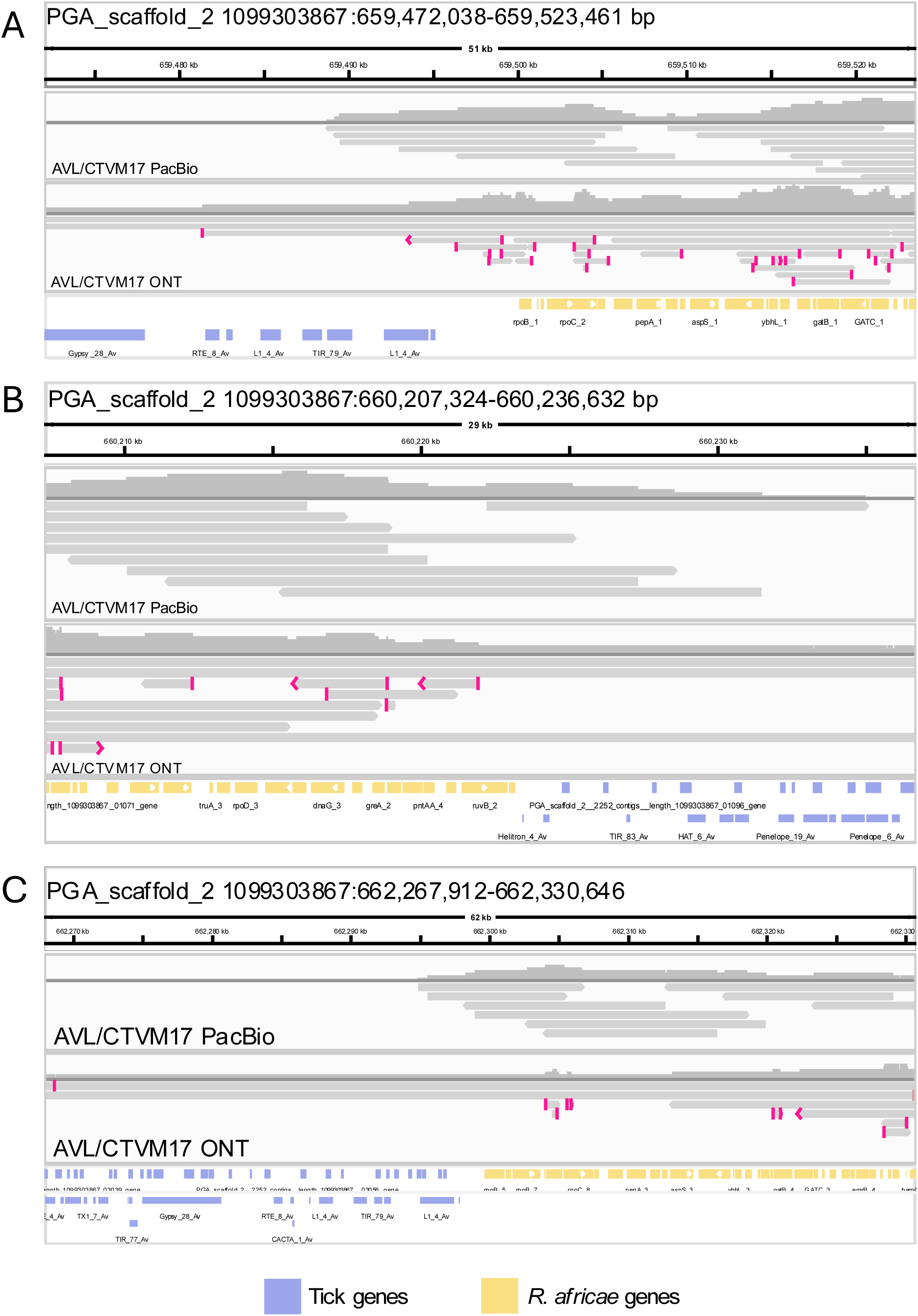
Mapping of AVL/CTVM17 PacBio and ONT libraries on PGA_Scaffold_2 of the Hi-C assembly. Sequence reads were observed spanning the junction between sequences containing tick and *R. africae* genes. Tick annotated regions flanking the insertion consist primarily of TEs.

**Fig. S6.**
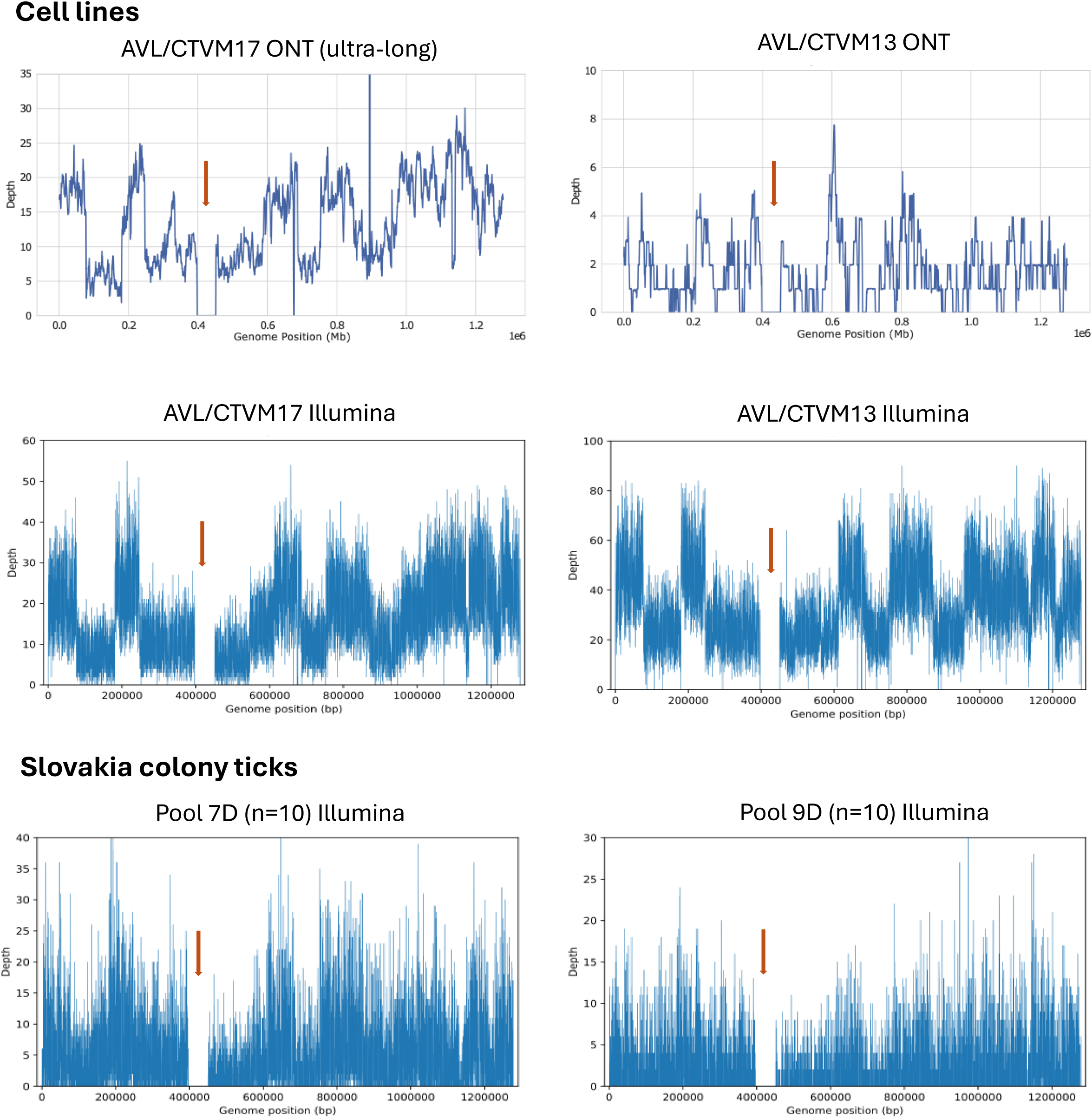
Coverage plots of sequence libraries from the indicated *A. variegatum* cell lines and Slovakia colony ticks mapped to the *R. africae* ESF-5 chromosome. Coverage plots were generated using mapped reads with 1000 bp rolling window. The arrows point to the missing 54 Kb region.

**Fig. S7.**
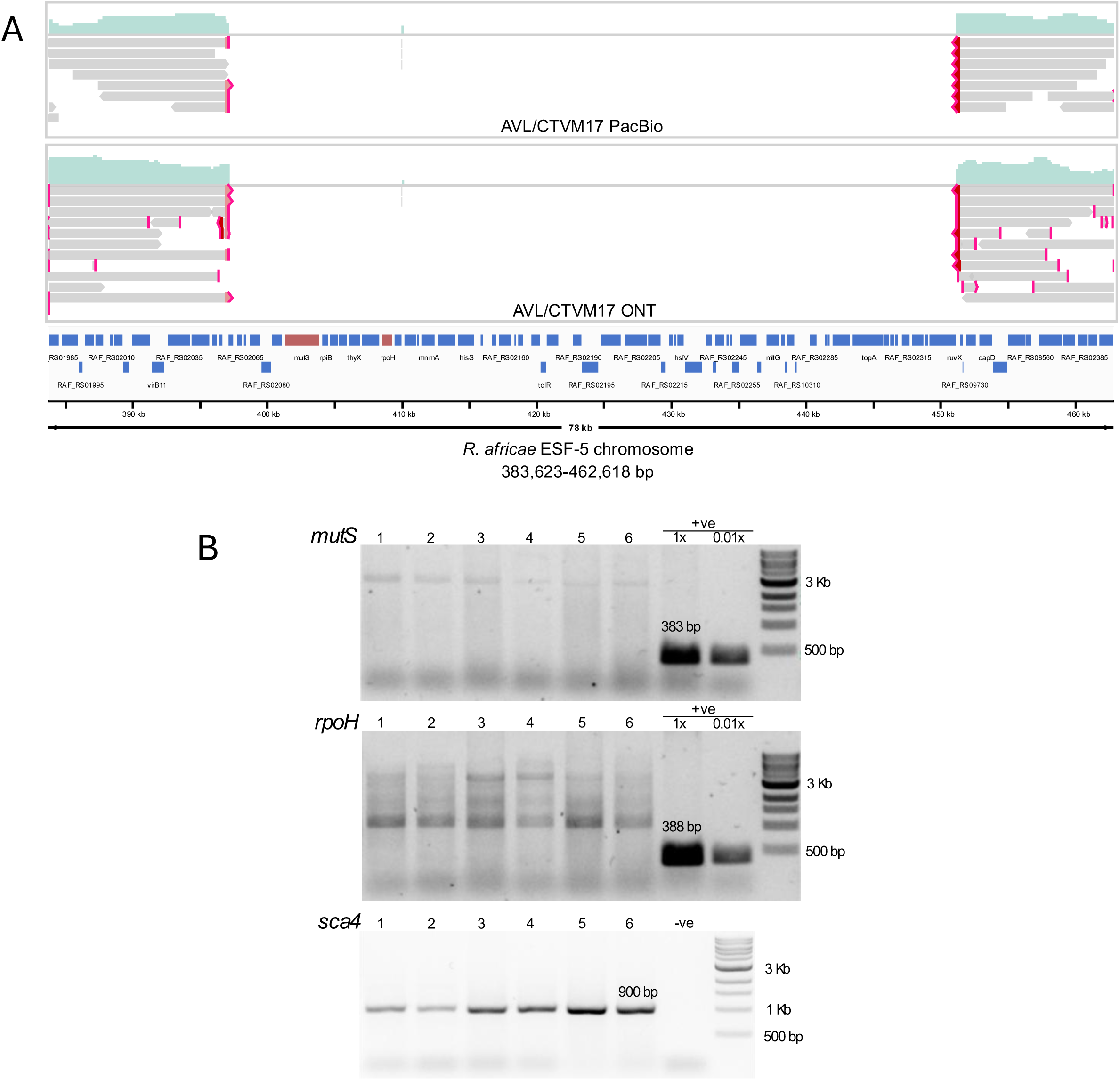
The 54 Kb missing region in *R. africae* insertion sequence. (A) Illumina sequences from *A. variegatum* AVL/CTVM17 cell line were aligned to the *R. africae* ESF-5 chromosome. The alignment was visualised using the Integrated Genomics Viewer browser. Coverage per nucleotide site (coloured in green) is presented above each alignment. Each grey horizontal bar represents a sequence read. Red ends and arrows on the bars indicate the rest of the sequence is aligned to other regions of the chromosome. Nucleotide substitutions, insertions and deletions are not displayed in the figure. Genes annotated on the *R. africae* chromosome are presented below the alignments. (B) The *mutS* and *rpoH* genes located in the missing region (coloured in red in A) were used as PCR targets to confirm the absence of these sequences in ticks from the Slovakian colony. Each lane represents DNA from an individual tick. The *sca4* PCR was performed as loading control – *sca4* is located in the insertion sequence. *R. africae* DNA extract from a clinical isolate was used as positive control.

**Fig. S8.**
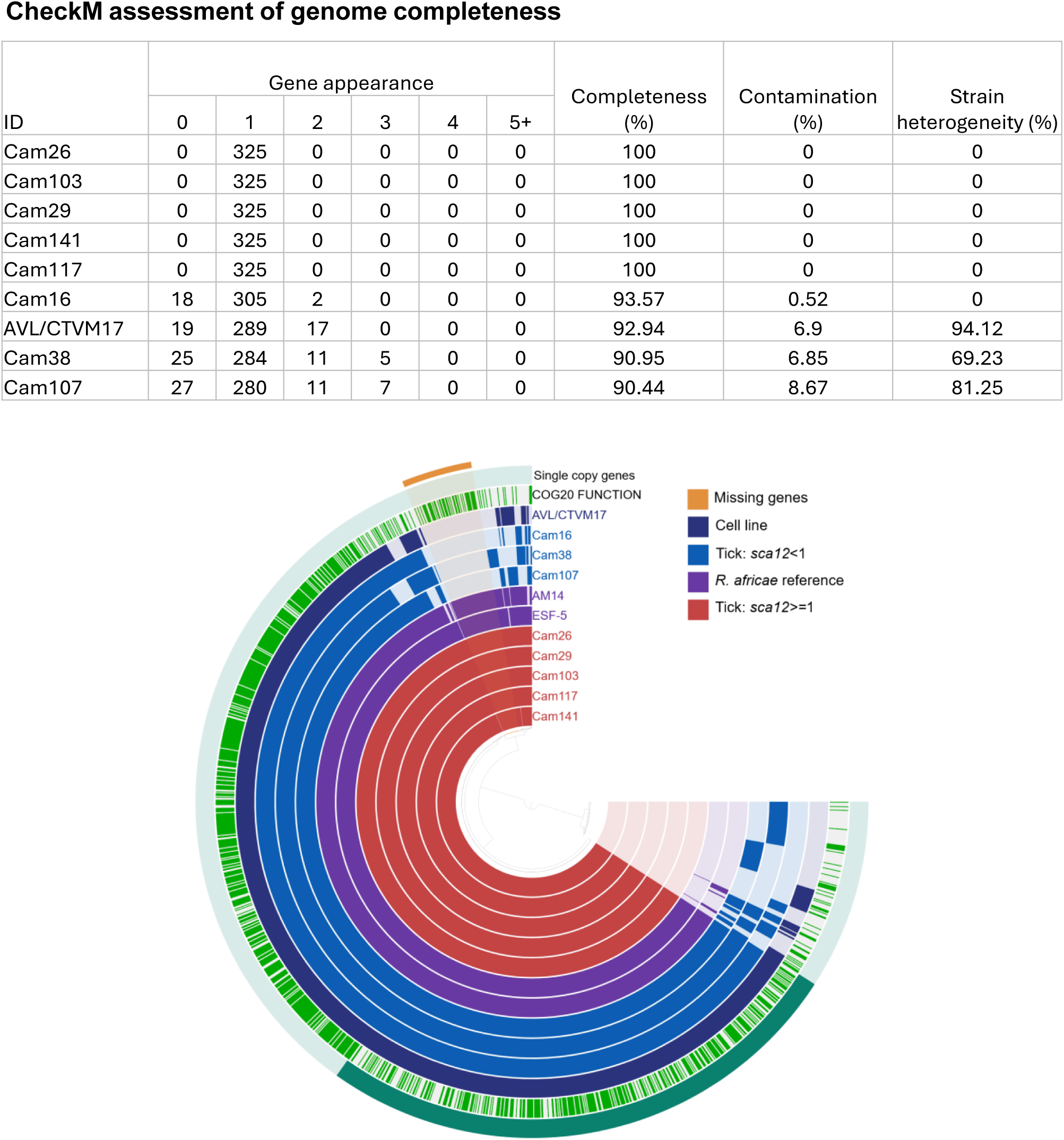
Pangenome analysis of *R. africae.* SPAdes assemblies were generated from the Illumina reads produced from field-collected ticks. The assembly of the *R. africae* insertion sequence in AVL/CTVM17 was generated with Flye v2.9 from PacBio and ONT reads. The SPAdes assembly of the *R. africae* AM14 was produced from publicly available Illumina data (NCBI SRA: SRX19868396). All sequence libraries were mapped to the *R. africae* ESF-5 chromosome sequence and the mapped reads were used for assembly. Contaminating sequences were removed from the assembly using Blobtools2. Genome completeness was assessed by CheckM using the Rickettsiales marker set (n=325, top panel). Pangenome was produced using the pangenomic workfIow in Anvi’o8 (bottom paneI). Each ring represents the presence (darker colour) or absence (lighter colour) of the gene clusters in the assembly.

**Table S6.**
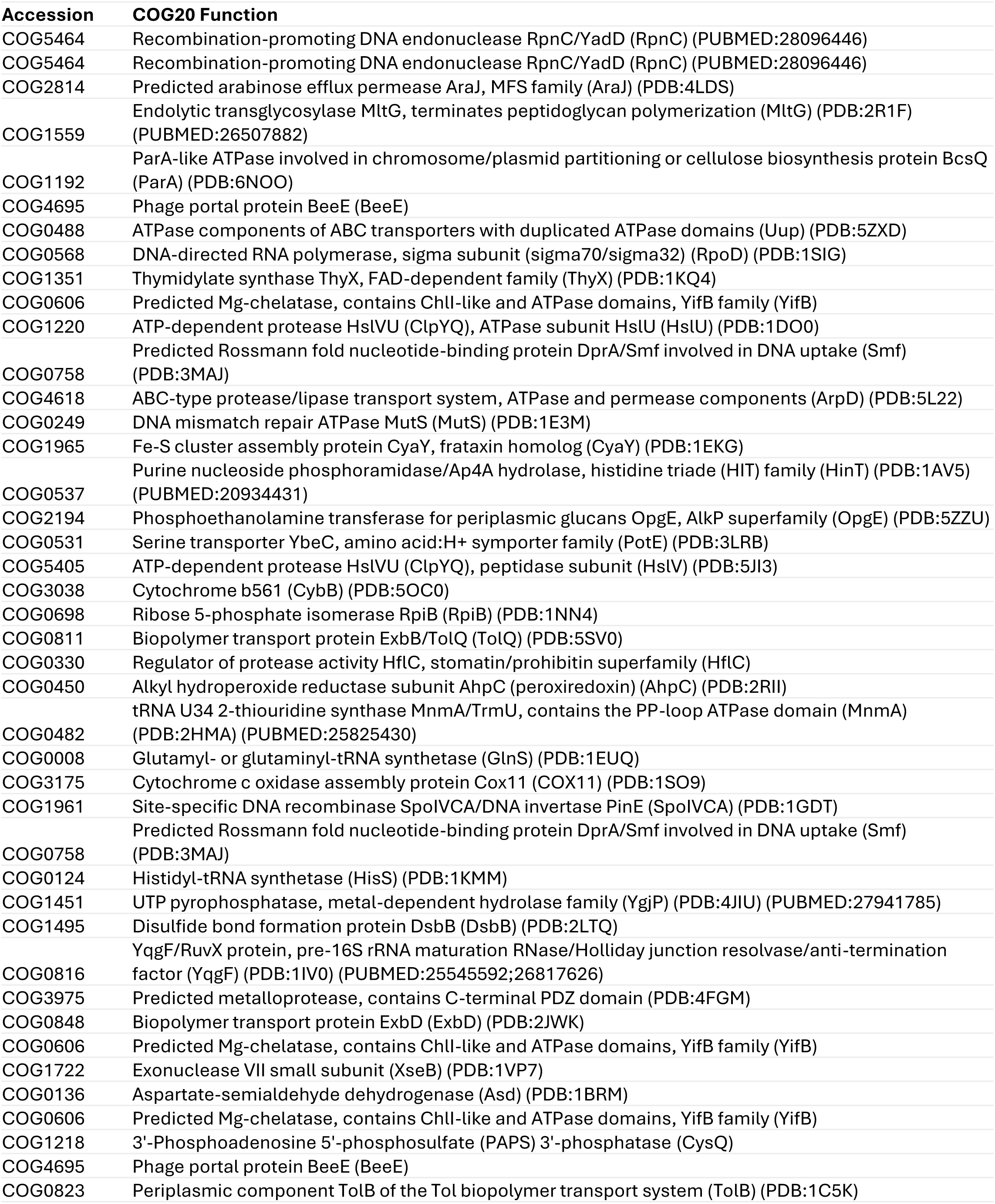
COG functions assigned to genes in the 54 Kb missing region.

