## Supplementary methods for "A near-complete chromosome from a bacterial pathogen integrated into the genome of its arthropod vector"

### 1. Antibiotic treatment assay

Cells of AVL/CTVM13 at passage (p) 148, AVL/CTVM17 cells at p120 and BME/CTVM23 cells at p78 were each seeded in 2-ml volumes into four replicate flat-sided culture tubes (Nunc). The BME/CTVM23 cells were then inoculated with supernate from a BME/CTVM23 culture heavily-infected with *Rickettsia raoultii*. Five days later (day 0), 200 µl of cell suspension was collected from each tube for DNA extraction. Two tubes from each cell line were then antibiotic-treated by addition of tetracycline hydrochloride (Sigma) to a final concentration of 100 µg/ml; the remaining two tubes of each cell line served as untreated controls. The cultures were then incubated at 32°C; medium was changed weekly by removal and replacement of ¾ of the medium volume, and fresh tetracycline was added weekly to each treated tube. Samples for DNA extraction were collected weekly up to day 35, from all cultures by pipetting the cells and removing 200 µl of cell suspension. Cell samples were centrifuged at 15,000 x g for 5 min, the supernate was discarded and the pellets were stored at – 20°C until the end of the experiment, when DNA was extracted from all samples at the same time using the DNeasy Blood and Tissue Kit (Qiagen, UK). The resultant DNA samples were subjected to quantitative PCR assays targeting fragments of the *Rickettsia* citrate synthase gene (*gltA*) and the tick *rpl6* gene described below to determine whether the tetracycline treatment resulted in any change in the proportion of the *Rickettsia gltA* gene, normalised to the tick housekeeping gene, in the treated versus untreated control cultures of each cell line.

### 2. Polymerase chain reactions (PCR)

DNA was extracted from aliquots of tick cell cultures or halves of field-collected ticks using the DNeasy Blood and Tissue Kit (Qiagen, UK). Rickettsial and tick qPCRs were performed using primers and protocols as previously described[1-3], except that primers for the tick-specific nuclear gene, *rpl6*, were redesigned to target metastriate ticks rather than *Ixodes* spp. [forward (5′-ACCCTACGGAAGACAGTCCA-3′) and reverse (5′- GTGATCCTCTTGCGGAGCTT-3′)]. Quantification of the *R. africae*-specific plasmid, pRa, was achieved using the *sca12* gene (unique to pRa) as the target with the primers (5′-CGTTAGAAGTCAAGGCGCAA-3′) and (5′-GAGCTGGTGGTAGTGACTGT-3′), probe (5′-TAAGGGCTCTGCA-GGAATTAGCACTAACGT-3′), and the same protocol as for the rickettsial *gltA* gene qPCR. Copy numbers obtained for *rpl6* were used to normalise the data for *gltA* and *sca12*.

For PCR amplification of the *mutS* gene, forward (5′- GTTAATGTACCGGAAACGGAAA-3′) and reverse (5′- TTTGTGACAATTTCAAGGTTGC-3′) primers were used; whereas PCR amplification of *rpoH* was achieved using forward (5′-GGAATGATTTTACAAGCTGCACA-3′) and reverse (5′-ACTTCATTAACCGAAACGCCTA-3′) primers. Both PCRs were performed with the following conditions: 3 min initiation at 95°C, 35 cycles of 1 min activation at 95°C, 30 s annealing at 50°C and 1 min elongation at 70°C, with a 7 min final elongation at 70°C. The PCR reactions were prepared in 20 µl containing 10 µl of 2X BioMix Red (Bioline, UK), 2 pmol forward and reverse primers, and 2 µl of DNA extract. The PCR for the *sca4* gene was performed as previously described[4]. A DNA extract from a clinical isolate of *R. africae* (strain PELE)[5], kindly provided by Prof. Marcelo Labruna, Universidade de São Paulo, Brazil, was used as the positive control.

#### 3. Fluorescence *in situ* hybridisation (FISH)

**Preparation of chromosome spreads**

To prepare metaphase chromosome spreads for FISH, a fresh culture of AVL/CTVM17 cells (p138) was seeded (1:1 split from a dense culture) on day 1. The next day, the culture was treated with colcemid (10 µg/ml, Roche Diagnostics, UK) and incubated overnight. On day 3, the cells were harvested by centrifugation (200 × g, 5 min) and resuspended in 0.6 % sodium citrate. The suspension was incubated at 37°C with occasional shaking for 35 min. The cell nuclei were then pelleted by centrifugation (400 × g, 5 min) and resuspended in ice-cold acetic alcohol (3 parts methanol, 1 part glacial acetic acid) for fixation. The fixation step was repeated once, and the pellet was resuspended in an equal volume of ice-cold acetic alcohol. For each microscope slide, 10 µl of the suspension was deposited onto the slide and warmed at 45°C on a heat block. A drop of 60% acetic acid was added onto the suspension. The cells in the suspension were then macerated and dispersed using a tungsten needle. The slide was transferred onto a frozen surface to continue with the maceration. The slide was left to air-dry and then dehydrated in an ethanol series (70%, 80%, 96%, each for 60 s).

**Probe hybridisation**

The air-dried slides were treated with 0.1 µg/µl RNase A (Invitrogen, UK) in 2x saline-sodium citrate (SSC) buffer (Invitrogen, UK) for 1 h at 37°C and washed three times with 2x SSC (Sigma-Aldrich, UK). A second fixation step was performed by incubating the slides in 3.7% formaldehyde diluted in 4X SSC, 0.1% v/v Tween 20 for 10 min. After washing three times with 2x SSC, the slides were dehydrated in an ethanol series (70%, 80%, 96%, each for 60 s). Biotin-labelled myTags custom probes were designed and manufactured by Daicel Arbor Biosciences, USA. These FISH probes (100 ng/slide) were prepared in hybridization solution (50% formamide, 10% dextran sulfate, 2x SSC) and denatured at 90°C for 10 min, and then chilled on ice for at least 5 min. The chilled probe mixture was then deposited onto the chromosome smear, covered with parafilm and incubated at 70°C for 1 min. The slides were immediately placed into a humidified chamber and incubated at 37°C in the dark overnight. The next day, the slides were incubated in 2x SSC at 42°C for 15 min followed by two washes with 2x SSC at room temperature for 5 min each and 0.2x SSC for 10 min.

**Fluorophore labelling**

Blocking of the chromosome spreads was performed at room temperate for 30 min using the ISH ReadyProbe blocking solution (Invitrogen, UK). The first layer of labelling was achieved by incubating the chromosome spread with an Alexa Fluor 488 streptavidin conjugate (Invitrogen, UK) at 1:1000 in blocking solution for 30 min at 37°C in a humidified chamber. The unbound fluorophore conjugates were then removed by two 5-min washes in 2X SSC/0.05% Tween 20 and a 5 min wash in 2X SSC. This was then followed by incubation with a biotinylated anti-streptavidin antibody (Vector Laboratories, UK) at 1:500 in ISH for 30 min at 37°C in a humidified chamber. Washing was performed as before to remove the unbound antibodies. The second layer of labelling was achieved by incubating the chromosome spread with Alexa Fluor 488 streptavidin conjugate and washing to remove the unbound conjugates as before. The chromosome spreads were then mounted in ProLong™ Diamond Antifade Mountant with DAPI (Invitrogen, UK) and covered with glass cover slips for preservation of the fluorescence signal.

**Confocal microscopy**

The chromosome spreads were visualised and imaged at the Centre for Cell Imaging (CCI), University of Liverpool, using a Zeiss Axio Observer.Z1/7 with LSM900 confocal microscope under the following settings:

**Objective:** Plan-Apochromat 63x/1.40 NA Oil objective (420782-9900-799)

| | |
|---|---|
| **Lasers:** | Ch1 – 488 nm wavelength, 0.2% power |
| | Ch2 – 405 nm wavelength, 1.0% power |
| **Detectors**: | Ch1 – GaAsP-PMT detector, range = 400 - 650nm, Gain = 620V |
| | Ch2 - GaAsP-PMT detector, range = 400 - 605nm, Gain = 650V |
| **Pinhole:** | Ch1 – 48 µm |
| | Ch2 – 42 µm |

**Acquisition Settings:** Pixel scan time = 0.73 µs, Averaging = 8, Nyquist sampling = 1x

For visualisation purposes, background fluorescence was cleared in ImageJ[6] by subtracting the average baseline noise calculated from five distinct cell-free extracellular regions. Measurements of the distance from the hybridisation signal to the furthest end of the chromosomes, as a proportion of the length of the entire chromosome, were performed using ImageJ[6].

### 4. Genomic DNA extraction, sequencing and assembly

**PacBio sequencing**

For PacBio sequencing, DNA was extracted from a single AVL/CTVM17 culture (p125) using the DNeasy Blood & Tissue kit (Qiagen, UK) according to the manufacturer's protocol. Library preparation and sequencing were performed at the Centre for Genomic Research at the University of Liverpool (CGR). TruSeq PCR-free paired-end libraries (2 x 150 bp) with a 350-bp insert were generated and sequenced on a HiSeq 4000. The CGR performed the following read curation: the raw fastq files were trimmed for the presence of Illumina adapter sequences using Cutadapt v1.2.1 with option -O 3[7]; the reads were further trimmed using Sickle v1.200 with a minimum window quality score of 20 (github.com/najoshi/sickle); and reads shorter than 20 bp after trimming were removed. The trimmed reads were assembled using hifiasm v.0.15.1[8]. Genome completeness was assessed using the Benchmarking Universal Single-Copy Orthologs (BUSCO) v5.2.2[9] with arachnida_odb10 dataset. Assembly of the mitochondrial genome from the PacBio dataset has been previously described[3].

**ONT sequencing**

Oxford Nanopore Technologies (ONT)-based sequencing was performed twice for AVL/CTVM17 cells (p125-127). The first run used the DNA preparation as above for library preparation according to the SQK-LSK109 protocol with a starting amount of 2 µg DNA and sequenced with MIN106 (R 9.4.1) flow-cells for MinION (ONT, UK), producing a 1D reads library. Raw fast5 files were base-called with ONT Guppy base-calling software v5.0.11 using dna_r9.4.1_450bps_sup configuration. Reads were analysed, filtered, trimmed and split according to quality using the Filtlong tool v.0.2.1 (github.com/rrwick/Filtlong).

The second run was performed at the University of Nottingham. Ultra-long DNA was extracted from AVL/CTVM17 cells using a previously published protocol[10]. The Ultra-Long DNA Sequencing Kit SQK-ULK001 was used for the library preparation and sequencing was done on GridION and PromethION (Oxford Nanopore Technologies, UK) devices at the University of Nottingham. The ONT libraries were then combined and used for genome assembly using Flye v2.8.2[11]. Sequencing library and assembly statistics were produced using QUAST v5.0.2[12].

**Illumina sequencing**

For Illumina sequencing, DNA was extracted from AVL/CTVM13 (p155) and AVL/CTVM17 (p139) cells, frozen salivary glands harvested from ten each of female and male Slovakia colony ticks, or from halves of individual field-collected Cameroonian ticks using the DNeasy Blood & Tissue kit (Qiagen, UK). Additionally, DNA extracts from ticks collected from southern Africa for a previous study[13] were also used for Illumina sequencing. Illumina libraries were prepared using the NEBNext Ultra II FS Kit (half-volume reactions) with a 350-bp insert and sequenced on a NovaSeq X Plus using 10B chemistry (paired-end, 2x150 bp) at the CGR. The CGR performed the following read curation: the raw fastq files were trimmed for the presence of Illumina adapter sequences using Cutadapt v4.5 with option -O 3[7]; the reads were further trimmed using Sickle v1.200 with a minimum window quality score of 20 (github.com/najoshi/sickle); reads shorter than 15 bp after trimming were removed; and only paired reads were retained.

5. **Hi-C sequencing and assembly**

Tissues from three male ticks from the Slovakia colony were ground in liquid nitrogen into a fine powder. Cells from a single AVL/CTVM17 culture (p130) were pelleted at 300 × g for 5 min. To generate cross-linked DNA, the fine tick powder and cell pellet were each resuspended in 1% formaldehyde solution and incubated at room temperature for 20 min. Equal volumes of 250 mM glycine were added to the suspensions and left to incubate for 15 min with occasional vortexing. The cross-linked material was pelleted, washed with sterile water and snap-frozen on dry ice before transporting to Phase Genomics (Seattle, WA, USA) for DNA extraction, Proximo™ Hi-C library preparation and sequencing on the Illumina S4 Novaseq platform. The BlobToolKit v4.2.1[14] was used to detect non-eukaryotic sequences. Scaffolding with the PacBio assembly was performed on Phase Genomics' Proximo™ genome scaffolding platform. Collinearity analysis was performed with MCScanX[15] to detect sequence duplications. A Circos plot was generated using Circa (https://circa.omgenomics.com). Mapping of PacBio and ONT sequence libraries to the Hi-C assembly was also inspected using the Integrative Genomics Viewer v2.17.4 (https://igv.org).

### 6. Table S1. Summary table for all materials sequenced in this study

European Nucleotide Archive sample accession numbers are provided for each sequence dataset used in the study.

| ***Sample*** | ***Source material*** | ***PacBio*** | ***Illumina*** | ***ONT-1D*** | ***ONT Ultra-long read*** | ***HiC*** | ***cDNA*** |
|---|---|---|---|---|---|---|---|
| *AVL/CTVM13* | Cell line | | ERS28453407 | ERS31033575 | | | |
| *AVL/CTVM17* | Cell line | ERS23954145 | ERS28453406 | ERS31033574 | SAMEA117512495 | ERS23954146 | ERS23954148 |
| *A. variegatum (n=3)* | Slovakia colony ticks (all tissues) | | | | | ERS23954147 | |
| *A. variegatum 7D (n=10)* | Slovakia colony ticks (salivary glands) | | ERS23954154 | | | | |
| *A. variegatum 9D (n=10)* | Slovakia colony ticks (salivary glands) | | ERS23954155 | | | | |
| *AML020* | Field-collected tick (halves) - Angola | | ERS28453408 | | | | |
| *AML069* | Field-collected tick (halves) - Angola | | ERS28453409 | | | | |
| *Cam16* | Field-collected tick (halves) - Cameroon | | ERS23954158 | | | | |
| *Cam103* | Field-collected tick (halves) - Cameroon | | ERS23954156 | | | | |
| *Cam107* | Field-collected tick (halves) - Cameroon | | ERS23954151 | | | | |
| *Cam117* | Field-collected tick (halves) - Cameroon | | ERS23954152 | | | | |
| *Cam141* | Field-collected tick (halves) - Cameroon | | ERS23954153 | | | | |
| *Cam166* | Field-collected tick (halves) - Cameroon | | ERS28453390 | | | | |
| *Cam168* | Field-collected tick (halves) - Cameroon | | ERS28453391 | | | | |
| *Cam176* | Field-collected tick (halves) - Cameroon | | ERS28453392 | | | | |
| *Cam178* | Field-collected tick (halves) - Cameroon | | ERS28453393 | | | | |
| *Cam183* | Field-collected tick (halves) - Cameroon | | ERS28453394 | | | | |
| *Cam184* | Field-collected tick (halves) - Cameroon | | ERS28453395 | | | | |
| *Cam196* | Field-collected tick (halves) - Cameroon | | ERS28453397 | | | | |
| *Cam205* | Field-collected tick (halves) - Cameroon | | ERS28453398 | | | | |
| *Cam210* | Field-collected tick (halves) - Cameroon | | ERS28453399 | | | | |
| *Cam221* | Field-collected tick (halves) - Cameroon | | ERS28453400 | | | | |

| | | |
|---|---|---|
| *Cam236* | Field-collected tick (halves) - Cameroon | ERS28453401 |
| *Cam239* | Field-collected tick (halves) - Cameroon | ERS28453402 |
| *Cam240* | Field-collected tick (halves) - Cameroon | ERS28453403 |
| *Cam247* | Field-collected tick (halves) - Cameroon | ERS28453404 |
| *Cam250* | Field-collected tick (halves) - Cameroon | ERS28453405 |
| *Cam26* | Field-collected tick (halves) - Cameroon | ERS23954157 |
| *Cam29* | Field-collected tick (halves) - Cameroon | ERS23954160 |
| *Cam38* | Field-collected tick (halves) - Cameroon | ERS23954150 |
| *MIMam052* | Field-collected tick (halves) - Mozambique | ERS28453419 |
| *MIMam144* | Field-collected tick (halves) - Mozambique | ERS28453418 |
| *MMCup011* | Field-collected tick (halves) - Mozambique | ERS28453420 |
| *MSMun030* | Field-collected tick (halves) - Mozambique | ERS28453415 |
| *MSMun063* | Field-collected tick (halves) - Mozambique | ERS28453416 |
| *MSMun316* | Field-collected tick (halves) - Mozambique | ERS28453417 |
| *ZiMG017* | Field-collected tick (halves) - Zimbabwe | ERS28453413 |
| *ZiMM007* | Field-collected tick (halves) - Zimbabwe | ERS28453414 |
| *ZLSY047* | Field-collected tick (halves) - Zambia | ERS28453410 |
| *ZML073* | Field-collected tick (halves) - Zambia | ERS28453411 |
| *ZPC002* | Field-collected tick (halves) - Zambia | ERS28453412 |

**7. Mapping of Illumina tick reads to the AVL/CTVM17 HiC Assembly**

Mapping of short reads from ticks to the ten largest AVL/CTVM17 Hi-C scaffolds was performed using bowtie2 v2.4.2 [7]. Coverage was determined using Mosdepth v0.3.11[16] at 100 Kb windows and extracted for each scaffold. Visualisation of read coverage from male and female ticks were performed in RStudio v2026.05.1+225 with the Tidyverse package v2.0.0.

**8. Phylogenetic tree inference**

BUSCO v5.2.2[9] was applied on each tick genome in Table S2, and the genome of the common house spider, *Parasteatoda tepidariorum*, as outgroup, to obtain single copy orthologue genes. Orthologous clusters containing a single gene copy from each species were identified with Orthofinder v2.5.4[17]. Each orthologous cluster was aligned with MAFFT v7.487[18] and trimmed with Gblocks 0.91b[19]. Alignments were concatenated and provided to model-test-ng v0.1.7[20] to assess the best suited model for tree inference. Finally, the alignment and model selection was provided to RaxML-ng v0.6.0[21]. The final tree was generated using the JTT+I+G4 model. RaxML-ng was performed using 10 randomised parsimony starting trees and 1000 bootstrap replicates.

**9. Table S2. Accession numbers for tick genome assemblies used in phylogenetic analysis**

| **Species** | **Accession** |
|---|---|
| *Amblyomma americanum* | GCA_030143305.2 |
| *Amblyomma maculatum* | GCA_023969395.1 |
| *Dermacentor andersoni* | GCF_023375885.1 |
| *Dermacentor silvarum* | GCF_013339745.2 |
| *Haemaphysalis longicornis* | GCA_013339765.2 |
| *Hyalomma asiaticum* | GCA_013339685.2 |
| *Ixodes ricinus* | GCA_043645445.1 |
| *Ixodes scapularis* | GCF_016920785.2 |
| *Rhipicephalus annulatus* | GCA_013436015.1 |
| *Rhipicephalus microplus* | GCA_013339725.1 |
| *Rhipicephalus sanguineus* | GCF_013339695.2 |
| *Parasteatoda tepidariorum* (outgroup) | GCF_000365465.3 |

#### 10. Genome annotation

The Hi-C assembly was first soft-masked using RepeatMasker v4.1.2[22] and a custom library was generated using Repeatmodeler v2.0.2[22]. The ONT data from cDNA sequencing was mapped to the genome using minimap2 v2.24[23] with the following parameters "-I 64G -t 32 -ax splice:hq -uf". A draft transcriptome was generated from mapped reads using stringtie2 v2.2.1[24]. Putative open-reading frames (ORFs) and untranslated regions were annotated using stringtie2 gtf output and TransDecoder v5.7.1 (https://github.com/TransDecoder/TransDecoder). TransDecoder also propagated the predicted ORFs back to the genome and provided a final set of coding sequences and protein sequences for each gene, as well as gene annotation in GFF3 format. Protein FASTA outputs were used for functional annotation using InterProScan v5.32-71.0[25], eggNOG v2.1.6[26] and SignalP v5.0b[27]. Putative functional annotations were applied to the GFF3 file using the AGAT toolkit[28]. Completeness of gene prediction was assessed using BUSCO v5.2.2[9].

#### 11. Gene family expansion analysis

For each tick species with public gene prediction sets, the longest isoform of each gene was extracted using the AGAT toolkit[28] and used for generating orthologous clusters using Orthofinder v2.5.4[17]. The Newick tree generated from above was trimmed to remove the nodes for *Amblyomma maculatum*, *Ixodes ricinus* and *Rhipicephalus annulatus*, as these did not have annotated protein sequences. The orthologous clusters and trimmed Newick tree were provided to CAFE5[29] for gene family contraction/expansion analysis, and the results were plotted with CafePlotter (https://github.com/moshi4/CafePlotter). The CAFE5 output and gene annotations from the above were parsed in RStudio v2024.04.2+764 (https://posit.co) to identify significantly expanded gene families. Terms related to mobile genetic elements, or TEs ("transposase", "integrase", "reverse transcriptase", "retrotransposon", "mobile element", "insertion sequence", "DDE", "Tnp" and "RVT") were used to search the Pfam annotations to identify the gene families related to TEs. To test if the expansion of TE clusters was unique to cell lines, the ratio of average Illumina read coverage per base for the identified TE genes to 50 randomly-selected single-copy genes was determined for the cell lines and selected ticks.

#### 10. TE library curation

The steps to obtain the curated TE library from the AVL/CTVM17 Hi-C assembly are illustrated in Figure SM1 and described as follows:

**Commented [LS1]:** For clarity, can you qualify this by stating what it's a library of - A. variegatum? AVL/CTVM17? Cell line and ticks?

1) RepeatModeler v2.0.3[30] was employed to predict *de novo* TEs using the LTRstruct option, generating 3,947 potential TE families.
2) Redundancy was eliminated using cd-hit-est (-d 0 -aS 0.8 -c 0.8 -G 0 -g 1 -b 500), resulting in 3,707 potential TE families.

3) Sequences shorter than 200 bp were eliminated, resulting in 3,146 potential TE families.
4) TE-Trimmer (https://github.com/qjiangzhao/TEtrimmer) was used to extend the consensus and eliminate false positives. This process identified 3338 potential TE families due to the separation of sequences in different subfamilies.
5) Additionally, for additional false-positive filtering and trimming of the long consensus created by TE-Trimmer, MCHelper v1.7.0 (https://github.com/GonzalezLab/MCHelper) was run with no extension step. Subsequently, 1539 potential TE families remained in the library.
6) Sequences identified as complete by MCHelper were manually inspected by analysing the TE-aid plots to confirm the presence of the basic structure of the transposable elements (TEs). Sequences lacking TE features were removed, and those containing non-TE edges or TE chimaeras were trimmed. From 760 potential TE families, 488 were confirmed.
7) The original RepeatModeler consensus sequences of the remaining potential TE were used to perform a blastn search against the 488 curated TE families (-outfmt "6 qseqid sseqid pident length mismatch qstart qend qlen slen" -evalue 1e-20 -reward 2 -penalty -3 -gapopen 5 -gapextend 2 -max_target_seqs 1). Sequences that matched an existing sequence in the library, with a minimum of 80% identity, were eliminated. Next, MCHelper v1.7.0 was used with three rounds of 1.5kb extension followed by manual inspection as described above. After this step, 236 TE families were confirmed.
8) An additional step was performed with the sequences that were eliminated by MCHelper due to the absence of complete copies in the genome. As TE-Trimmer can eventually produce an unrealistic consensus, we retrieved the original RepeatModeler consensus sequences, performed MCHelper with one round of 1.5 kb extension, and eliminated sequences that matched existing TEs in the library as described above. After manual inspection, 137 additional sequences were confirmed.
9) As a last step, to ensure real TEs were not eliminated during the curation steps, the presence of the original RepeatModeler sequences in the curated TE library was investigated by BLASTn as described above. The potentially missing TE families were then inspected for the presence of TE domains using PFAM[31]. A total of 104 sequences were rescued in this step.
10) After blast searches of known tick TEs missing in the library, three additional families were added: Ruka, a SINE element previously identified in ixodid ticks[32]; the DNA transposon *Merlin* identified in *Ixodes scapularis*[33]; and a SINE element similar to SINE2-2_DeSi from *Dermacentor silvarum* (as identified in online Repeat Masking in Repbase).
11) The final curated TE library contains 965 sequences that were either classified during the manual inspection step or using the RepeatClassifier module[30]. Non-autonomous LTRs were classified in their respective superfamilies if they contained at least a partial coding region allowing the classification. The remaining non-autonomous LTRs were considered as LARD (if larger than 2.2 kb) or TRIM only if the direct terminal repeats were clearly identified as LTRs with the diagnostic starting nucleotides TG.

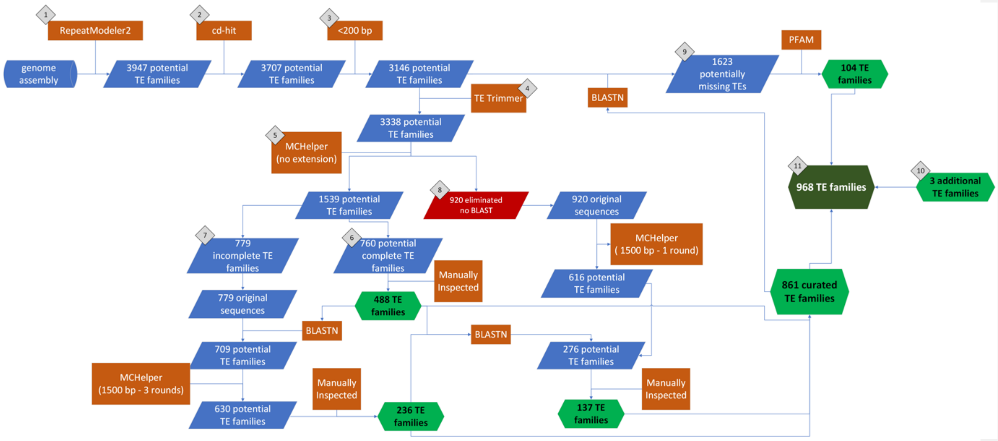

Figure SM1. Workflow for the identification and curation of the TE library. This workflow combined automated tools with manual inspection steps to validate and classify TE families. The number of TE families identified and retained is indicated. Details of each step in the workflow are provided in the main supplementary text.

### 11. *Rickettsia africae* and nuclear mitochondrial DNA (NUMT) insertion sequence analysis

The *R. africae* ESF-5 reference and *A. variegatum* mitochondrial (Accession: PZ344169.1) assemblies were aligned to the AVL/CTVM17 Hi-C scaffolds using lastz v1.04.22[34] and visualised using JBrowse2 v/cli/2.10.3[35]. Mapping of short reads from ticks to the AVL/CTVM17 Hi-C scaffolds was performed using bowtie2 v2.4.2[7]. Coverage was determined using Mosdepth v0.3.11[16] with 100-Kb windows. Coverage across the entire PGA_scaffold_2 or only to the region with the *R. africae* insertion (659.6–664.2 Mb) was extracted for comparison. Statistical tests and data visualisation were performed in RStudio v2026.05.1+225 with the Tidyverse package v2.0.0 (https://www.tidyverse.org). Wilcoxon tests were performed for comparisons between groups, assuming a non-normal distribution as determined from Shapiro-Wilk normality tests. The indicated *A. variegatum* cell line assemblies and Illumina reads from Slovakia colony tick were also mapped to the *R. africae* ESF-5 chromosome sequence. Mapped contigs/reads were extracted using SAMtools (v1.19.2) *view*[36] and coverage was determined using Mosdepth v0.3.11[16]. A Python script was used to plot the coverage across the *R. africae* ESF-5 chromosome sequence with a 1-Kb rolling window.

Mapping of short reads from ticks and AVL/CTVM17 to the *R. africae* ESF-5 reference assembly was performed using bowtie2 v2.4.2[7]. Coverage was determined using Mosdepth v0.3.11[16] with 1-Kb windows and visualised in RStudio. Unpaired t-tests were performed for comparison of the means, assuming normal distribution as determined from Shapiro-Wilk normality tests.

**Commented [JK2]:** I assume the mitochondria paper will be out first.

For single nucleotide polymorphisms (SNPs) analysis, the resulting BAM files were randomly down-sampled to 50X coverage in order to minimise potential biases arising from uneven sequencing depth. Subsequently, SNPs were called using Freebayes v1.3.8[37]. The resulting vcf files were sorted using BCFtools v1.17[36] and filtered using VCFtools v0.1.17[38] with the following parameters: --remove-indels --minQ 100 --recode --recode-INFO-all. Principal component analysis was performed using PLINK2[39] with the following parameters: --double-id --allow-extra-chr --maf 0.01 --freq. The resulting principal components were visualised in RStudio using the ggplot2 package. The analysis was performed three times with different down-sampling seed values (42, 101 and 202) to assess consistency.

#### 12. ***Rickettsia africae*** **genome assembly and pangenome analysis**

The PacBio and ONT reads from the AVL/CTVM17 cell line were combined and mapped to the *R. africae* ESF-5 reference assembly using minimap2 v2.28-r1209[23]. The mapped reads were used for assembly of the *R. africae* insertion genome using metaFlye v2.9.3-b1797[40]. Short reads from ticks in this study and *Amblyomma sparsum* AM14 (Genbank SRA: SRR24067184)[41] mapped to the *R. africae* ESF-5 reference assembly were used to assemble the genome for *R. africae* using the SPAdes genome assembler v3.15.5[42]. The BlobToolKit v4.2.1[14] was used to remove eukaryotic contaminants from the assembly. Gene calling and annotation were performed using Prokka v1.14.5[43]. Genome completion was assessed using CheckM v1.2.2[18] and pangenome analysis was conducted using the pangenome workflow in Anvio v8 [44].

#### 13. ONT cDNA reads analysis

The ONT cDNA reads from AVL/CTVM17 were mapped to PGA_scaffold_2 of the Hi-C assembly using minimap2. Mapped reads were extracted using Samtools *view*. Gene features and mapped reads in BAM format were passed to bedtools[45] for average coverage-per-gene calculation. The output TSV file was used for plotting in R studio. Bedtools was also used to calculate average coverage in 10-Kb windows across the insertion region, as well as 1 Mb either side (658 Mb to 665 Mb). The output TSV file was used for plotting in R studio.
